# Myofibroblast- specific autophagy drives cyst growth in autosomal dominant polycystic kidney disease

**DOI:** 10.64898/2026.06.05.730503

**Authors:** Abeda Jamadar, Viji Remadevi, Meekha M Varghese, Haichun Yang, Vedant P. Thakkar, Indra Chandrasekar, Wen-Xing Ding, Volker H. Haase, Darren P. Wallace, Reena Rao

## Abstract

**Background:** Autosomal dominant polycystic kidney disease (ADPKD) is characterized by progressive cyst expansion, fibrosis and inflammation, leading to kidney failure. Myofibroblasts (MFs) often accumulate around cysts and promote fibrosis and cyst growth, but the cellular mechanisms enabling their pro-cystogenic activity remain unclear. Here we examined the role of autophagy within MFs, on their paracrine stimulation of cyst expansion in ADPKD.

**Methods:** Autophagy was assessed in human ADPKD nephrectomy tissue, primary human ADPKD renal myofibroblasts (ADPKD-MFs) and male RC/RC mouse model of ADPKD using immunostaining, LC3/p62 analyses, and transmission electron microscopy. Autophagy in MFs was inhibited pharmacologically in ADPKD-MFs, or by conditional Atg5 deletion in PDGFRβ-expressing renal stromal cells in RC/RC (RC/RC;Atg5KO) and wild type (WT;Atg5KO) mice.

**Results:** In human and mouse ADPKD kidneys, we detected LC3 puncta and autophagic organelles within αSMA-expressing MFs. Inhibition of autophagy in ADPKD-MFs blocked their paracrine stimulation of cyst epithelial cell proliferation *in vitro*. RC/RC;Atg5KO mice showed significantly reduced cystic growth, fibrosis, MF abundance, and improved kidney function. WT;Atg5KO mice showed no abnormalities in kidney structure or function. Targeted metabolomics performed on ADPKD cyst epithelial-cell conditioned media (ADPKD-ECs CM) revealed moderate increase in lactate levels compared to normal human kidney epithelial-cell conditioned media. Furthermore, lactate treatment stabilized hypoxia-inducible factor-1α (HIF1α) in myofibroblasts, while pharmacological inhibition of HIF1α reduced the expression of autophagy-related genes and impaired autophagic flux.

**Conclusion:** These findings reveal that autophagy in MFs is a previously unrecognized driver of cyst expansion and fibrosis in ADPKD. Lactate-mediated HIF1α stabilization in MFs promotes autophagy that is required for their paracrine stimulation of cyst epithelial growth. Targeting MF-specific autophagy or its upstream regulators may represent a therapeutic strategy to limit cyst growth and fibrosis in ADPKD.

## Background

ADPKD is the most common hereditary cause of kidney disease in the United States(1), and affects more than 12.5 million individuals worldwide (2) . A defining feature of ADPKD is the development of fluid-filled and epithelial cell lined cysts originating from kidney tubules. This process is accompanied by progressive fibrosis and inflammation that lead to the ultimate loss of kidney function (3–5). The cyst microenvironment in ADPKD kidneys often includes dense and remodeled extracellular matrix (ECM), myofibroblasts (MFs) and inflammatory cells. The MFs are typically α-smooth muscle actin (αSMA) expressing cells which play an essential role in wound repair, and are the primary producers of ECM in chronic diseases of most organs, including the kidneys (6–10) and in tumors (11).

Large populations of MFs accumulate in ADPKD kidneys, frequently surrounding cysts (12, 13). We previously identified a pathogenic reciprocal interaction between MFs and the cyst-lining epithelium in ADPKD kidneys (12, 13). Cyst epithelial cell-secreted factors stimulate MF activation and accumulation in the cyst microenvironment (13), and MFs in turn, directly stimulate cyst growth (12). Depletion of MFs (12), or treatment with the FDA approved antifibrotic drugs, Nintedanib (14) or Pirfenidone (15) in ADPKD mouse models not only reduced kidney fibrosis but also inhibited cyst epithelial cell proliferation and cyst growth. In this study, we sought to define the role of autophagy in renal interstitial MFs, in their paracrine regulation of cyst growth and ADPKD progression.

Autophagy is a pro-survival catabolic process activated by nutrient depletion and hypoxia (16), enabling cells to degrade intracellular proteins and recycle amino acids to sustain protein synthesis and tricarboxylic acid cycle activity (16–18). Beyond metabolic support, autophagy maintains cellular homeostasis by clearing damaged organelles and misfolded proteins through the formation of autophagosomes, which fuse with lysosomes to form autolysosomes for enzymatic degradation (19).

In ADPKD kidneys, cyst epithelial cells exhibit markedly reduced autophagy, a defect implicated in disease progression (20, 21). However, the role of autophagy in ADPKD remains controversial, as re-activation of autophagy has been reported to either suppress (22) or promote (23) cyst growth in vertebrate models of the disease. These conflicting observations may reflect cell type-specific autophagy functions within the heterogeneous cellular environment of the kidney. In the tumor microenvironment, autophagy in stromal cells is known to support tumor growth, disease progression, and metastasis (24). Here we sought to investigate the role of autophagy in renal myofibroblasts, and its contribution in ADPKD pathogenesis. We found that autophagy in MFs plays an important role in regulating cyst epithelial growth, and that autophagy in ADPKD MFs can be induced in a HIF1α-dependent manner by metabolites such as lactate, secreted from cyst epithelial cells.

## Materials and Methods

### Reagents

Sodium L-lactate (#L7022) and Chloroquine diphosphate salt (#C6628) from Sigma Aldrich, Burlington, MA, USA., Bafilomycin A1 (#HY100558) from MedChemExpress, Monmouth Junction, NJ, USA, and LW6 (#6322) and DMOG (#4408) from Tocris Bioscience (Biotechne, MN, USA) were used.

### Human tissue samples and primary cells

Kidney tissue and cells from normal human kidneys (NHK) and ADPKD kidneys were de-identified samples. Primary culture MFs (ADPKD_MFs) were isolated using a protocol modified from a previously published method (25). Briefly, fibrotic tissue explants (10-15 mm³) were dissected from human ADPKD nephrectomy kidneys and minced using a scalpel and forceps under sterile condtions. Tissues were enzymatically digested for 2 hours in 1% collagenase type II solution (#LS004176, Worthington Biochemical Corp., Lakewood, NJ, USA), followed by gently dissociation by pipetting and passaged through a 70 µm cell strainer to remove undigested tissue. To inactivate collagenase, complete medium (DMEM/F12, 10% FBS and 1% Pen/Strep) was added to the filtrate in a 2:1 ratio, and the suspension was centrifuged at 2000 rpm for 2 minutes. The resulting cell pellet was resuspended in fresh complete medium and seeded onto collagen-coated (#354236, Corning, NY, USA) 100 mm petri dishes. After 10 mins incubation at 37°C, the cell suspension was removed, and the petri dishes were washed with complete media twice to remove non-adherent cells. The cell suspension was then transferred onto a new set of collagen-coated 100 mm petri dishes followed by incubation at 37^0^C for 10 mins to repeat this method of differential adhesion. The adherent cells were incubated in complete medium and at 37^0^C with 95% O_2_ and 5% CO_2_ until the cells reached 90% confluency, which typically took around 8-10 days. Myofibroblasts were identified by positive immunostaining for aSMA and vimentin and negative for pan-cytokeratin, an epithelial cell marker (Supplementary Fig 1A). QRTPCR analysis showed significant increase in aSMA, FSP and vimentin mRNA levels in ADPKD MFs compared to ADPKD epithelial cells (Supplementary Figure 1B). Flow cytometric analysis demonstrated that 97% of ADPKD myofibroblasts expressed Thy-1 (CD90), confirming a highly enriched fibroblast population (Supplementary Figure 1C). Normal human kidney (NHK) fibroblasts were purchased from Cell biologics (#H-6016, Chicago, IL) and used as controls.

### Targeted Metabolomics analysis

Monolayers of human primary culture NHK and ADPKD kidney epithelial cells were grown in 100 mm petri dishes. To collect cell culture conditioned media (CM), confluent monolayers of cells were incubated in serum free media, and media was collected after 24 hours. The CM was centrifuged at 2000 rpm for 5 minutes to remove cell debris and supernatant stored at -80 °C until further analysis. Media was thawed on ice and 50 µL-100 µL aliquots were pipetted for organic acid analysis, respectively. Media aliquots for organic acids were spiked with 13C isotopically-labelled organic acid internal standards, derivatized with ortho-benzylhydroxylamine, extracted in ethylacetate, and the extracts were dried under nitrogen at 45 °C according to previously published methods (26). The extracts were reconstituted in 50% methanol and quantitated using similarly prepared standard calibration curves with an Agilent 1290 Infinity II HPLC/6495B triple quadrupole mass spectrometer (26). Media aliquots for amino acids spiked with 13C isotopically-labelled organic acid internal standards, derivatized with a Waters Accq-Tag derivatization kit, and quantitated using similarly prepared standard calibration curves with an Agilent 1290 Infinity II HPLC/6495B triple quadrupole mass spectrometer (26).

### Mouse models

The Pkd1^RC/RC^ mouse (RC/RC) carrying a p.R3277C mutation is an orthologous model of ADPKD (27). A tamoxifen inducible platelet derived growth factor-β (PDGFRβ) *Pdgfrβ-P2A-CreER^T2^* was used to delete *Atg5* in the RC/RC mice. To obtain RC/RC;Atg5KO mice, RC/+ mice were bred with *Atg5*^f/f^ mice and *Pdgfrβ-P2A-CreER^T2^* mice (B6.Cg-*Pdgfrb^tm1.1(cre/ERT2)Csln^*/J, Strain# 030201, Jackson Laboratory, Bar Harbor, ME) (28). At 4 months of age, aSMA expressing myofibroblasts were detected in the RC/RC mouse kidneys, and some of them showed LC3 puncta (Supplementary Fig 2A). To induce *Atg5* gene knockout, mice were treated with tamoxifen (#T2859, Sigma Aldrich, Burlington, MA, USA) at a dose of 200 mg/Kg, by intraperitoneal injection. Mice were injected once a month, at 4, 5 and 6 months of age to induce recombination in the newly appearing MFs. To control for any potential effects of tamoxifen on cyst growth, all mice were administered tamoxifen. Wild type (WT) (n=8), Atg5KO (n=8), RC/RC (n=9) and RC/RC;Atg5KO (n=9), littermate mice were sacrificed at 6.5 months of age. Male littermate mice of mixed background were used in the study. Mice from each litter were assigned to study groups as they reached 4 months of age. No randomization was used. The investigators who injected the mice, quantitated cysts, cell proliferation and fibrosis, measured blood urea nitrogen (BUN) levels were blinded to the genetic identity of the mice. Mice were housed in a temperature-controlled environment, and experimental mice were placed on the same rack. All mice were sacrificed between 11:30 AM and 12:30 PM. No adverse effects were observed. No mice or datapoints were excluded. All mouse related studies were approved by the University of Kansas IACUC committee. ARRIVE guidelines were followed for animal studies (29). At sacrifice, blood was collected and plasma isolated. Kidneys were weighed and flash frozen or fixed in 4% paraformaldehyde.

### Picrosirius red staining

Paraffin embedded mouse kidney tissue sections were stained with Picrosirius red to visualize collagen fibers following manufacturer’s protocol (ScyTek Laboratories, #PSR-2, Logan, UT, USA). Polarized light microscopy and image analysis software (Qupath) were used for the quantitative assessment of kidney fibrosis. The amount of collagen deposition was quantified by measuring the percentage of area of birefringence-positive staining relative to the total kidney area.

### BUN

Plasma BUN levels were measured using QuantiChrom Urea Assay Kit (DIUR-100, BioAssay Systems, Hayward, CA, USA) as per the manufacturer’s protocol.

### Statistics

Values are expressed as mean±SEM for *in vivo* and mean±SD for *in vitro* studies. Data was analyzed by two-tailed unpaired T-test with Welch’s correction; or one-way ANOVA followed by Tukey’s multiple comparison test using GraphPad Prism software (Version 10). P≤0.05 was considered significant.

### Supplemental methods

Detailed methods for collection of CM, *in vitro* BrdU incorporation (30), Annexin V-FITC/PI staining (31), transmission electron microscopy (32), cyst quantification (33), Western blot (34), immunohistochemistry/ immunofluorescence staining (35), super resolution microscopy-vibratome sectioning and immunostaining protocol (36) and quantitative real-time PCR (33) are provided in Supplementary methods. The list of primer sequences for QRTPCR is provided in Supplementary Table 1.

## Results

### Renal interstitial myofibroblasts in ADPKD kidneys undergo autophagy

To determine whether ADPKD renal myofibroblasts (ADPKD-MFs) undergo autophagy, we examined microtubule-associated protein light chain 3 (LC3), a marker for autophagosomes. During autophagic flux, LC3-I is lipidated to LC3-II and incorporated into autophagosome membranes (37). Examination of human ADPKD nephrectomy kidneys and RC/RC mouse kidneys showed LC3 granular puncta in the αSMA expressing MFs (Fig 1A, B and Supplementary Fig 2B, C and Supplementary video-1). Transmission electron microscopy confirmed autophagic organelles in interstitial cells of RC/RC kidneys (Supplementary Fig 3A, B, C). LC3 puncta were also detected in primary culture human ADPKD renal MFs (Fig 1C). These findings indicate that ADPKD-MFs in both human and mouse kidneys exhibit autophagy.

**Figure 1:**
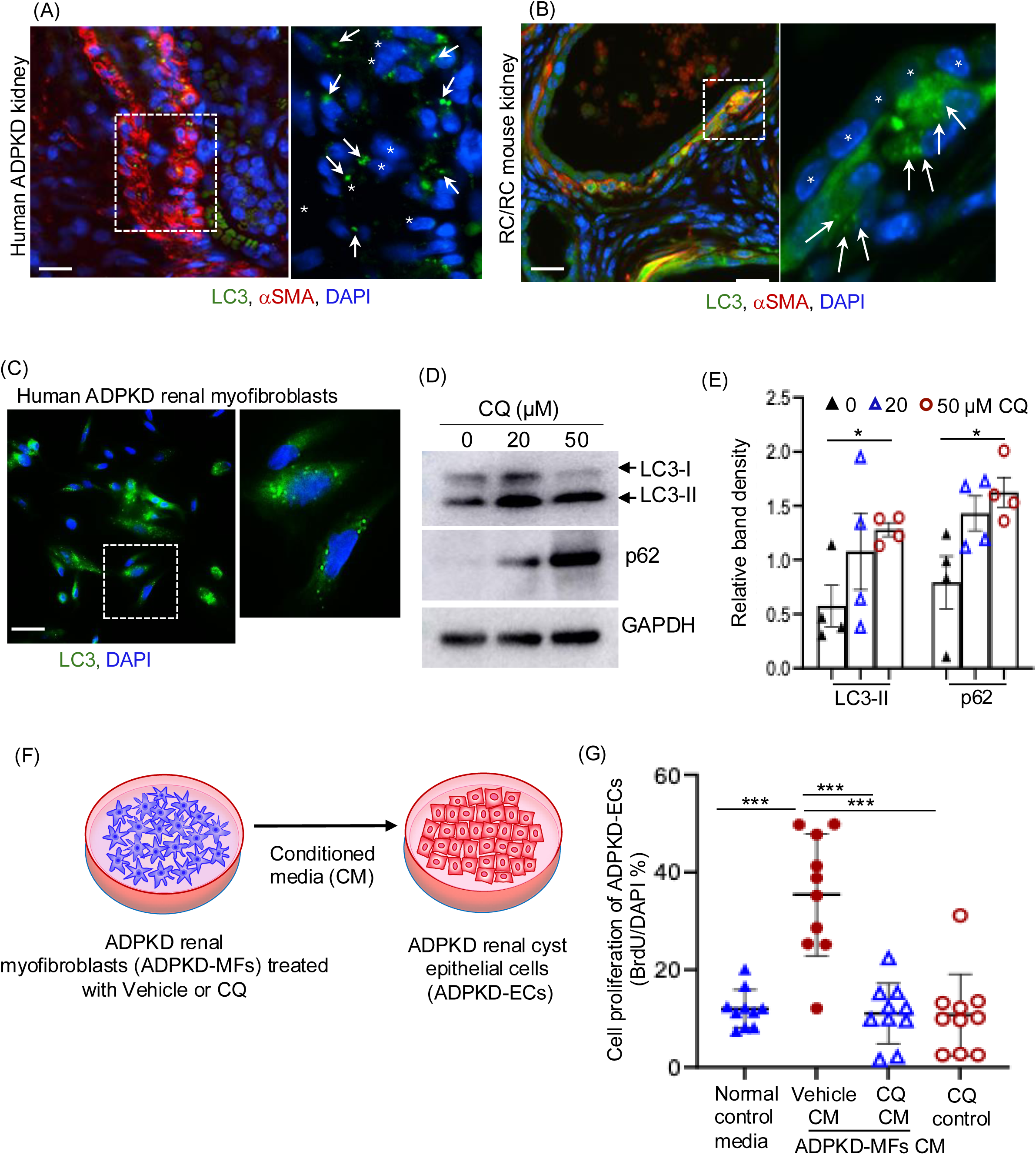
Myofibroblasts (MFs) undergo autophagy in human and mouse ADPKD kidneys, which is important to stimulate ADPKD renal cyst epithelial cell (ADPKD-ECs) proliferation. (A) Immunostaining for αSMA (red, denotes MFs), LC3 (green, denotes autophagosome) and DAPI (blue, denotes nuclei) in human ADPKD kidney, and (B) mouse kidney tissue sections of 6.5 months old RC/RC mice. The arrows points to LC3 puncta, and *denotes cyst lining epithelial cells (Scale bar = 200µm). (C) LC3 puncta in cultured ADPKD-MFs grown for 24h in serum free media (Scale bar = 200µm). (D) Immunoblot of ADPKD-MFs treated with 20µM and 50µM chloroquine (CQ) for 24 hours. (E) Quantitation of band density for Fig 1D. (F & G) ADPKD renal epithelial cells (ADPKD-ECs) were exposed to cell culture conditioned media (CM) from ADPKD-MFs treated with vehicle or CQ (20µM for 24h), or normal control media (not exposed to any cells), or directly treated with CQ (control) for a total 24h. *P<0.05 by T-test for E and ***P<0.001 by one-way ANOVA for G.

### Autophagy in ADPKD myofibroblasts is required for paracrine stimulation of cyst epithelial cell proliferation

To determine if autophagy in MFs supports cyst epithelial cell proliferation, we inhibited autophagy in ADPKD-MFs using chloroquine (CQ) *in vitro*. During effective autophagic flux, the cargo protein p62, which tags ubiquitinated aggregates to autophagosomes undergoes lysosomal degradation. Thus, impaired autophagic flux results in accumulation of both LC3-II and p62 (38). CQ treatment increased LC3-II and p62 in ADPKD-MFs, indicating autophagy inhibition (Fig 1D,E Supplementary Fig 4). Human ADPKD renal cyst epithelial cells (ADPKD-ECs) exposed to cell culture conditioned media (CM) from vehicle-treated ADPKD-MFs (Fig 1F) showed increased cell proliferation compared to cells cultured in normal control media not exposed to any cells (Fig 1G). However, CM from CQ treated ADPKD-MFs failed to increase cell proliferation of ADPKD-ECs (Fig 1G). Direct CQ treatment of ADPKD-ECs did not alter their proliferation (Fig. 1G). These findings indicate that autophagy in ADPKD-MFs is required for their paracrine promotion of cyst epithelial cell proliferation.

### *Atg5* gene knockout in PDGFRβ+stromal compartment reduces cyst growth in RC/RC mouse kidneys

To test the role of autophagy in MFs in cyst growth, we generated RC/RC mice with inducible deletion of *Atg5*, a key autophagy gene (39) using *Pdgfrβ-P2A-CreER^T2^*. In kidney fibrosis, PDGFRβ marks mesangial cells, interstitial fibroblasts and pericytes (40), all of which are capable of differentiating into myofibroblasts (6). Only male mice were used in this study, as previous finding indicate no significant sex-dependent differences in cyst progression in this model (41). All mice were sacrificed at 6.5 months of age. RC/RC;*Atg5*^f/f^;*Pdgfrβ-P2A-CreER^T2^* (RC/RC;Atg5KO) mice were compared with wildtype (WT), *Atg5*^f/f^;*Pdgfrβ-P2A-CreER^T2^* (Atg5KO) and RC/RC littermates (Fig 2A). We found increased LC3-I and p62 levels in RC/RC;Atg5KO kidneys compared to RC/RC kidneys (Supplementary Fig 5A, B), consistent with autophagy inhibition, and similar to findings from tissue-specific Atg5 gene deletion in mouse liver (42).

**Figure 2:**
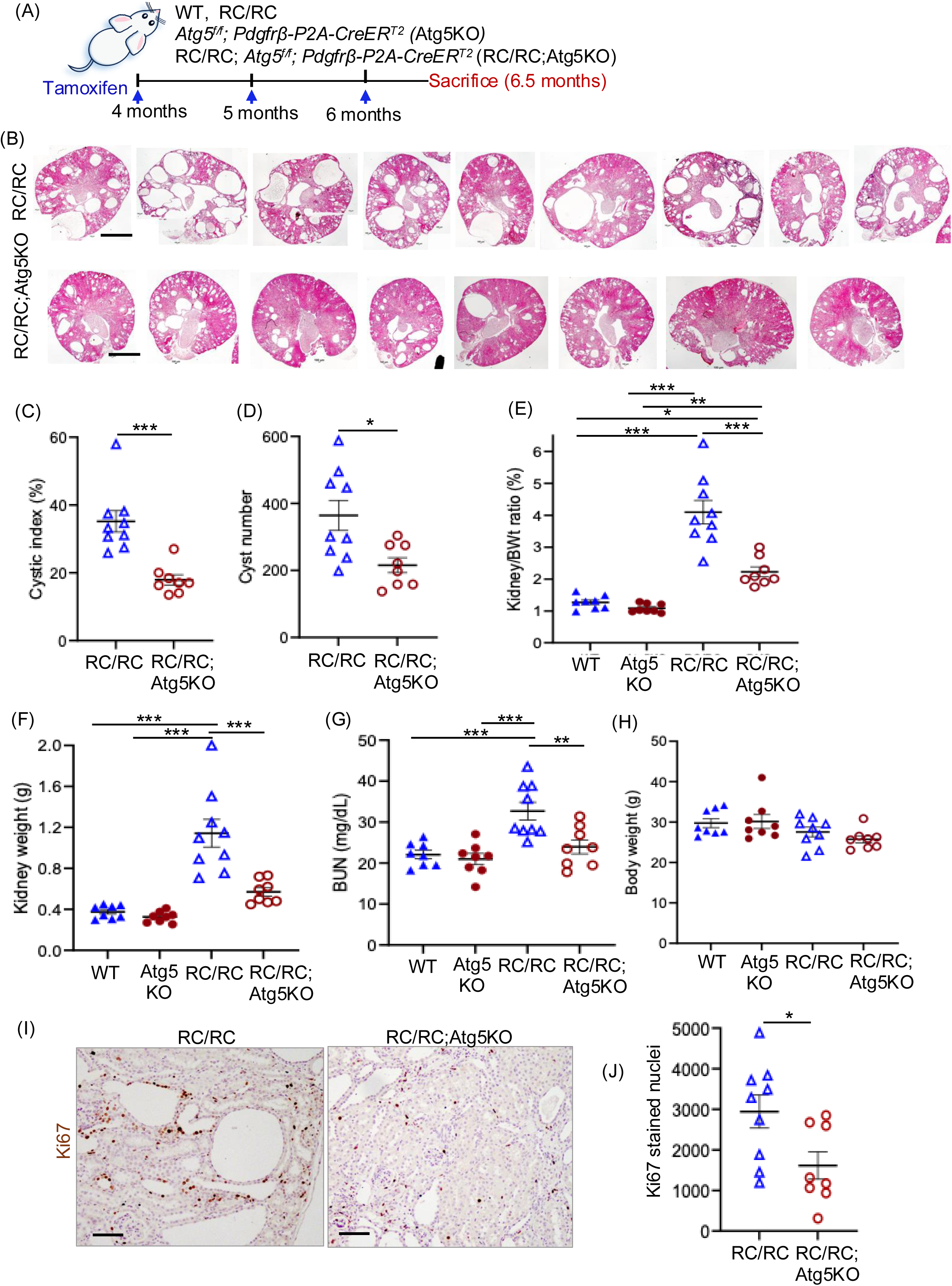
*Atg5* gene knockout in PDGFRb+stromal compartment reduces cyst growth in RC/RC mouse kidneys: (A) Male WT, Atg5KO, RC/RC and RC/RC;Atg5KO male were injected three times with tamoxifen (one dose of 200 mg/Kg/ month) at 4, 5 and 6 months of age and all the mice were sacrificed at 6.5 months of age. (B) H&E-stained kidney tissue sections (Scale bar = 1mm). Each kidney section is taken from the individual mice. (C) Cystic index (% cyst area/ total area of the kidney per tissue section), (D) Cyst number, (E) Two kidneys to body weight ratio, (F) Total kidney weight, (G) Blood urea nitrogen (BUN). (H) Body weight. (I) Immunostaining for Ki67 (brown nuclear staining) in mouse kidney tissue sections (Scale bar = 100µm) and (J) Quantitation of Ki67 staining per kidney section. Each data point represents individual mice. *P<0.05, **P<0.01, ***P<0.001 by T-test for C, D and J and *P<0.05, **P<0.01, ***P<0.001 by one-way ANOVA for E, F and G.

The RC/RC;Atg5KO mice showed reduced cyst growth as shown by H&E staining of kidney tissue sections (Fig 2B), decreased cystic index (Fig 2C) and cyst number (Fig 2D). The kidney to bodyweight ratio (Fig 2E), kidney weight (Fig 2F) and BUN levels (Fig 2G) in the RC/RC;Atg5KO mice were significantly reduced when compared to RC/RC mice. The Atg5KO mice on WT background showed no significant difference in kidney to body weight ratio, kidney weight or BUN compared to WT control mice (Fig 2E,F,G). No change in the bodyweight was observed across all the groups (Fig 2H). Moreover, Atg5KO mice did not exhibit any renal morphological abnormalities compared with WT mice (Supplementary Fig 5C). The RC/RC;Atg5KO kidneys also showed ∼50% reduction in cell proliferation by Ki-67-staining compared to RC/RC kidneys (Fig 2I,J). These results show that *Atg5* gene deletion in PDGFRβ-expressing cells reduces cyst growth and improves kidney function in the RC/RC mouse model of ADPKD.

### *Atg5* gene knockout in PDGFRβ+stromal compartment attenuates MFs expansion and kidney fibrosis

To determine the impact of autophagy inhibition on kidney MFs, we measured αSMA expression. The RC/RC;Atg5KO kidneys showed significantly reduced αSMA protein levels by Western blot (Fig 3A,B) and immunostaining (Fig 3C), as well as decreased αSMA (*Acta2*) mRNA levels (Fig 3D), indicating a smaller MF population compared to RC/RC kidneys. Examination of cleaved caspase-3 revealed no significant differences between RC/RC and RC/RC;Ag5KO kidneys as assessed by western blot (Supplementary Fig 6A, B) and immunostaining (Supplementary Fig 6C, D), suggesting no change in apoptotic cell death. *In vitro*, CQ-treated ADPKD-MFs similarly showed no increase in apoptosis as determined by Annexin V/ propidium iodide (PI) staining using flow cytometry (Supplementary Fig 6E).

**Figure 3:**
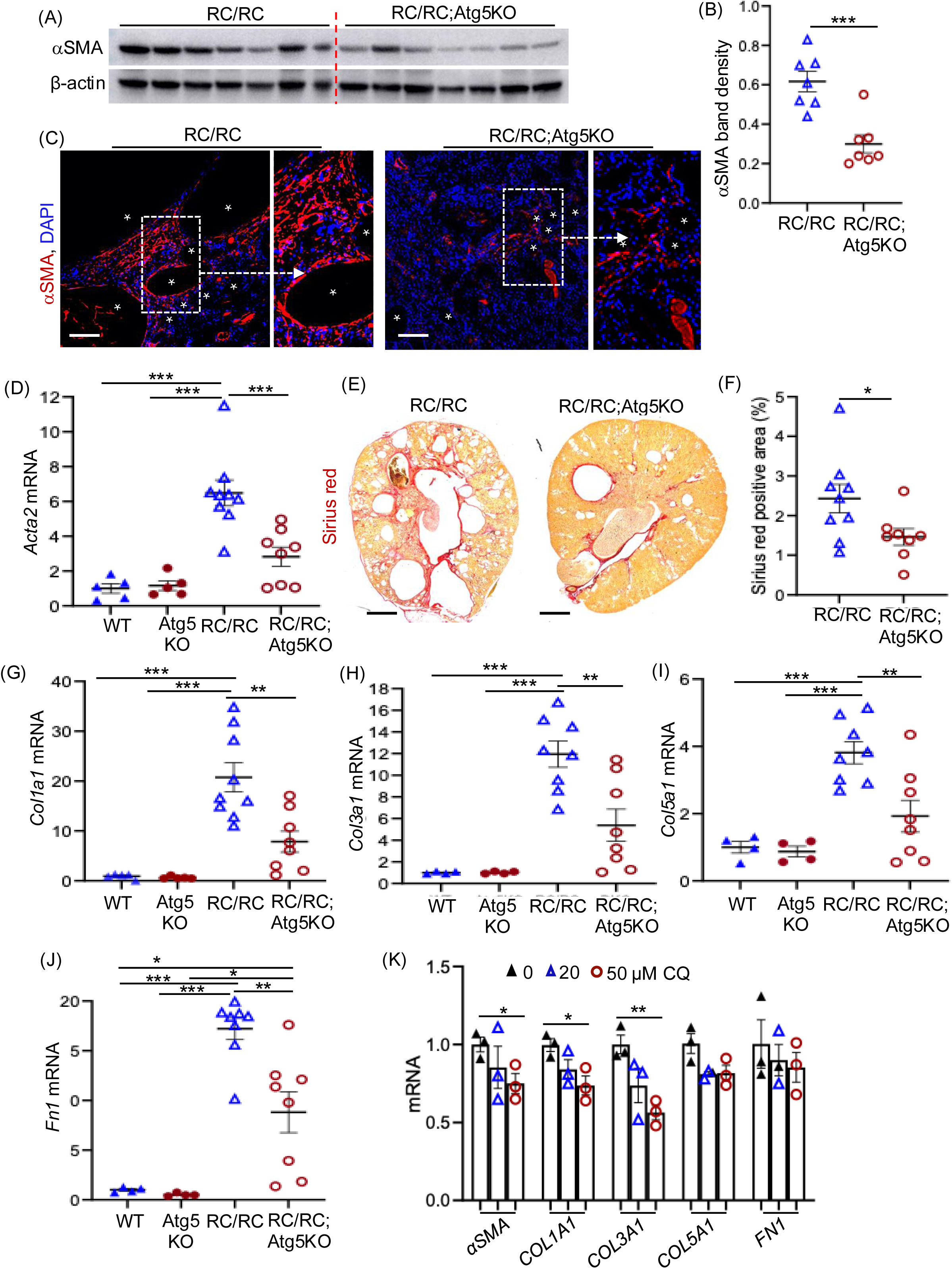
*Atg5* gene knockout in PDGFRb+stromal compartment reduced kidney fibrosis in RC/RC mouse kidneys: (A) Immunoblot on kidney tissue lysate and (B) quantitation of band density for αSMA (C) Immunostaining for αSMA staining (red), DAPI (blue) and *denotes cyst (Scale bar - 100 µM). (D) QRTPCR for *Acta2* in mouse kidney tissue lysate (E) Picrosirius red staining in mouse kidney tissue sections (Scale bar 1mm). (F) Quantitation of Picrosirius red staining by polarized light microscopy. QRTPCR for (αSMA), (G) *Col1a1* (Collagen 1a1) (H) *Col3a1* (Collagen 3a1) (I) *Col5a1* (Collagen 5a1) and (J) *Fn1* (Fibronectin) in mouse kidney tissue lysate. Each data point represents individual mice. (K) QRTPCR for ECM genes in ADPKD MFs treated with 20 and 50 mM CQ for 24 hours. *P<0.05, ***P<0.001, T-test for B, F and K and *P<0.05 **P<0.01, ***P<0.001 by one-way ANOVA for D, G, H, I & J.

RC/RC;Atg5KO kidneys also displayed reduced fibrosis, with lower type-1 collagen, indicated by Sirius red staining and quantitation (Fig 3 E,F), and decreased mRNA levels of collagens *Col1a1*, *Col3a1*, *Col5a1*, and fibronectin (*Fn1*) compared to RC/RC kidneys (Fig 3G-J). Moreover, inhibition of autophagy by chloroquine treatment (50 µM) in human ADPKD-MFs significantly reduced the mRNA expression levels of *aSMA*, *COL1A1* and *COL3A1* compared to control-treated cells, whereas *COL5A1* and *FN1* did not show a significant reduction (Figure 4K). Overall, these results demonstrate that autophagy suppression in PDGFRβ-expressing cells limits myofibroblast expansion and attenuates kidney fibrosis in an ADPKD mouse model.

**Fig 4.**
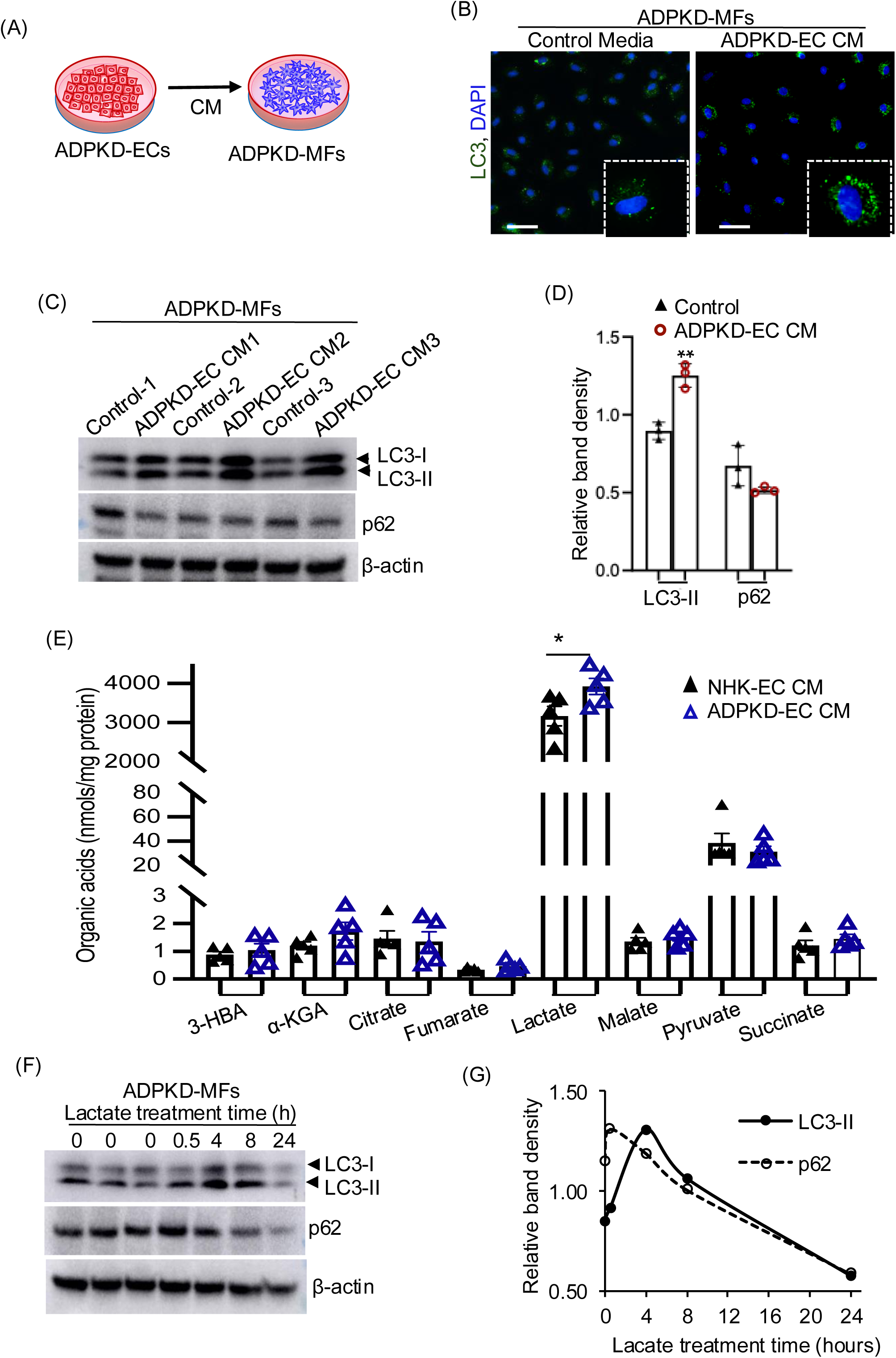
ADPKD-ECs regulate autophagy in interstitial MFs in ADPKD kidneys: (A) Cartoon illustrating ADPKD MFs exposed to cell culture conditioned media collected from ADPKD ECs (B) ADPKD-MFs exposed to ADPKD-ECs CM or normal control media (not exposed to any cells) for 24 hours were immunostained for LC3 (green) and DAPI (blue) in MFs (Scale bar = 200µm). (C) Cell lysates were immunoblotted for LC3-I, LC3-II and p62 and (D) Quantitation of band density for Fig 4B. (E) Metabolomic analysis of ADPKD-ECs and normal human kidney (NHK)-ECs CM (n=5 human kidneys in each group). Each “n” represents a biological replicate (human ADPKD kidney samples). (F) Immunoblot for ADPKD-MFs treated with lactate (20 mM) at different timepoint (G) quantitation of band density for Fig 4E. *P<0.05, **P<0.01 by T-test for D and E.

### Lactate secreted from cyst epithelial cells induces autophagy in ADPKD-MFs

To determine whether ADPKD-ECs promote autophagy in ADPKD-MFs, we exposed ADPKD-MFs to ADPKD-ECs CM (Fig 4A). ADPKD-MFs showed baseline LC3 puncta after 24 hours in serum-free control media which was not exposed to any cells (Fig 4B). Exposure to ADPKD-ECs CM further increased LC3 puncta in ADPKD-MFs (Fig 4B), elevated LC3-II levels, compared to control media, indicating enhanced autophagic flux (Fig. 4C,D). To identify components of ADPKD-ECs CM that induce autophagy in MFs, we performed targeted metabolomics for organic acids in ADPKD-ECs CM and normal human kidney epithelial cell CM (NHK-ECs CM) (Fig 4E, Supplementary Excel File 1). Lactate levels were increased by 20% in ADPKD-ECs CM when compared to NHK-ECs CM, whereas other organic acids showed no significant differences (Fig 4E). Direct treatment of ADPKD-MFs with 20 mM lactate increased LC3-II levels, reduced p62 levels (Fig 4F, G) and increased LC3 puncta (Supplementary Fig 7). These results suggest that lactate, which is elevated in ADPKD-ECs CM, could be a paracrine factor capable of inducing autophagy in ADPKD-MFs.

The oxygen-regulated α-subunit of HIF-1 (HIF1α) is expressed in cyst-lining epithelial cells and contributes to PKD pathogenesis (43), but its role in kidney MFs is unclear. Given that HIF-1 regulates autophagy in cancer and liver disease (44, 45), we examined whether it influences autophagy in ADPKD-MFs. Immunostaining showed elevated HIF1α expression and increased nuclear localization in ADPKD-MFs compared with Normal Human Kidney (NHK) fibroblasts (Fig 5A). HIF1α levels increased further when ADPKD-MFs were exposed to lactate or ADPKD-ECs CM (Fig 5B,C, Supplemental Fig 8A,B), consistent with prior studies showing lactate-mediated stabilization of HIF1α through inhibition of prolyl hydroxylases (PHD) (46).

**Figure 5:**
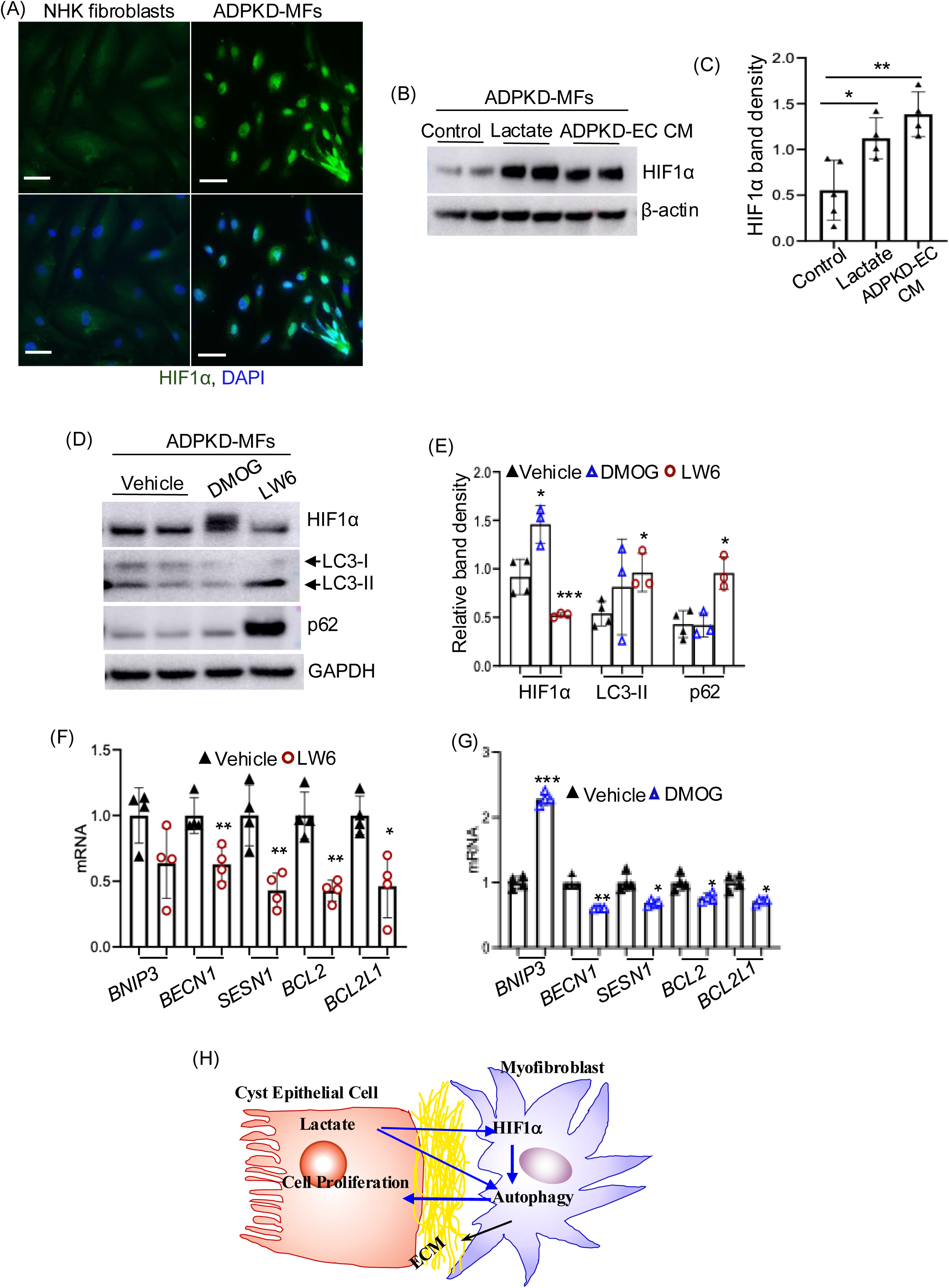
HIF1α regulates autophagy in ADPKD-MFs: (A) Immunostaining of HIF1α (green) and DAPI (blue) in normal human kidney fibroblasts and ADPKD-MFs (Scale bar = 200µm). (B) Immunoblot for ADPKD-MFs treated with lactate (20 mM) or ADPKD-ECs CM for 8 hours and (C) its quantitation of band density for Fig 4G and supplementary Fig 6A, B. (D) Immunoblot for ADPKD MFs treated with DMOG (1mM) or LW6 (20 µM) for 24 hours and its (E) Quantitation of band density for Fig 4I and supplementary Fig 6C. (F) mRNA expression of autophagy related HIF1α genes in ADPKD-MFs treated with vehicle or LW6 (20 µM) for 16 h. (G) mRNA expression of autophagy related HIF1α genes in ADPKD-MFs treated with vehicle or DMOG (1mM) for 16 h. (H) Cartoon demonstrating cyst epithelial-MFs interaction. Each “n” represents a biological replicate (human ADPKD kidney samples). *P<0.05, **P<0.01, ***P<0.001 by T-test for C, E, F and G.

Under normoxia, the canonical oxygen-sensing pathway is active, where PHDs hydroxylate HIF1α leading to proteasomal degradation of HIF1α. To examine the role of HIF1 in autophagy in MFs, we used dimethyloxalylglycine (DMOG), a pan-PHD inhibitor to stabilize HIF1α, and LW6 which promotes PHD-independent HIF1α degradation (47). LW6 treatment reduced HIF1α expression, whereas DMOG increased HIF1α expression relative to vehicle treatment in ADPKD-MFs (Fig 5D,E, Supplementary Fig 8C). Notably, LW6 increased p62 levels and caused LC3-II accumulation, indicating impaired autophagy upon HIF1α inhibition (Fig 5D,E, Supplementary Fig 8C). In contrast, DMOG did not further alter p62 or LC3-II compared with vehicle (Fig 5D,E, Supplemental-Fig 8C). Analysis of autophagy-related genes showed that LW6 decreased Beclin1, Sestrin1, BCL2 and BCL-XL (Fig 5F), while DMOG increased BNIP3 and decreased Beclin1, Sestrin1, BCL2, and BCL-XL in ADPKD-MFs (Fig 5G). Together, these findings suggest that autophagy in ADPKD-MFs is supported in part by lactate derived from cyst-lining epithelial cells and by HIF1α accumulation in MFs (Fig 5H).

## Discussion

Our studies show that the MFs in human ADPKD and RC/RC mouse kidneys exhibit active autophagy. Autophagy in ADPKD-MFs supports cyst growth because, its inhibition (A) by chloroquine reduced cyst epithelial cell proliferation *in vitro*, and (B) by conditional knockout of *Atg5* in PDGFRb+stromal cells in RC/RC mice reduced kidney cyst growth and fibrosis, and improved kidney function. Mechanistically, ADPKD cyst epithelial CM, rich in lactate, induced autophagy in MFs *via* stabilization of HIF1α. Inhibition of HIF1α impaired autophagy and downregulated autophagy-related gene expression in MFs. These findings show that autophagy in MFs is important for cyst growth, and that a lactate-HIF1α mediated mechanism could regulate it.

Our study identifies a novel role for autophagy in kidney MFs within the pathogenesis of ADPKD. While autophagy maintains cellular homeostasis under stress and is implicated in diverse physiological and pathological processes (48, 49), its role in cystogenesis has been controversial. In ADPKD, cyst-lining epithelial cells show impaired autophagy (21). Enhancing autophagy *via* Beclin-1 upregulation suppressed cyst development, while autophagy inhibition by systemic *Atg5* gene deletion exacerbated cystogenesis in a Pkd1-deficient zebrafish model (22). In contrast, autophagy activation promoted cyst growth in mice and zebrafish models of PKD (23). Further complicating the picture, treatment of PKD mice with trehalose, an autophagy-inducing disaccharide, showed no significant effect on cyst progression (50, 51). We found that MFs surrounding cysts undergo autophagy in human ADPKD kidneys and RC/RC mouse kidneys, as evidenced by LC3 puncta and autophagic organelles detected *via* immunohistochemistry and electron microscopy. Human ADPKD-MFs also displayed autophagic activity *in vitro*, supporting the relevance of these findings across species. Critically, inhibition of autophagy in MFs blocked their paracrine stimulation of cyst epithelial cell proliferation, indicating that autophagy in MFs is not merely a stress response but actively drives cyst growth.

To directly test the *in vivo* relevance of autophagy in MFs, we deleted Atg5 in the PDGFRb+stromal cells of RC/RC mice. This led to reduced kidney size, cyst burden, epithelial cell proliferation, and improved kidney function. Notably, the RC/RC;Atg5KO kidneys showed reduced αSMA and ECM expression, indicating that autophagy in MFs supports not only cyst growth but also fibrosis. These findings align with prior evidence that autophagy is critical for fibroblast activation in cancer and heart disease (52–55); persistent autophagy promotes kidney fibrosis, whereas autophagy inhibition attenuates it in unilateral ureteral obstruction (39). Similarly, cancer-associated fibroblasts use autophagy to drive tumor progression, and its inhibition enhances immunotherapy efficacy (56). Given that MFs-derived ECM and interstitial remodeling can support cyst expansion in ADPKD (3, 57–59), the observed reduction of ECM could also have indirectly contributed to reduced cyst growth in RC/RC-Atg5KO mice.

Our studies show paracrine interaction between MFs and epithelial cells in ADPKD kidneys. While macrophages and T-cells are known to influence cyst growth in ADPKD (60–64), we previously showed that ADPKD cyst epithelial cells stimulate MFs proliferation, differentiation, and migration (13), and conversely MFs directly stimulate cyst growth by paracrine signaling (12). Moreover, epithelial - mesenchymal cell paracrine crosstalk has been implicated in cyst expansion in PKD (65, 66).

In the current study, we found that elevated lactate in ADPKD-ECs CM induces autophagy in MFs. ADPKD epithelial cells and Pkd1^-/-^ mouse embryonic fibroblasts exhibit high glucose uptake and lactate production, hallmarks of enhanced aerobic glycolysis (67, 68). Lactate can enhance autophagy *via* lactylation of VPS34, a catalytic subunit of the class III PI3K complex, in cancer cells and skeletal muscle (69). Thus, lactate functions not only as a metabolic byproduct, but also as a paracrine regulator of autophagy in ADPKD-MFs. However, ADPKD-ECs CM likely contains additional factors that modulate autophagy. For example, CCN1 and CCN2 induce autophagy in cardiomyocytes (70) and osteoblasts (71) respectively, and we have shown that CCN2 secreted by ADPKD cyst epithelial cells promotes MF accumulation and fibrosis (12). Therefore, the observed increase in autophagy in MFs cannot be solely attributed to lactate in epithelial cell-CM.

We found that ADPKD-ECs CM, or lactate alone are sufficient to stabilize HIF1α in human ADPKD-MFs. We therefore asked whether HIF1 activation itself can enhance autophagic flux in MFs and contribute to ADPKD progression. HIF1α expression is elevated in ADPKD kidneys, and studies in rodent PKD models show that tubular HIF1α inactivation suppresses cyst growth, whereas pharmacological stabilization of HIF1α accelerates cyst expansion (43, 72–74). The HIF pathway also contributes to cystogenesis in other genetic diseases, including von Hippel-Lindau disease, hereditary leiomyomatosis and renal cell carcinoma syndrome. In ADPKD, excessive ECM deposition and interstitial remodeling, together with elevated luminal and interstitial fluid pressures, compress and distort kidney architecture, limiting oxygen and nutrient delivery to cyst-lining epithelial cells. (75–78). While hypoxia provides a physiological stimulus for HIF activation in the ADPKD kidneys (79), our *in vitro* studies show that HIF activation in MFs can occur under normoxia. Our finding that pharmacological reduction of HIF1α by LW6 impaired autophagic flux in ADPKD-MFs, indicates that HIF1α is at least partly required for lactate-induced autophagy. While LW6 promotes VHL-mediated proteasomal degradation of HIF1α even under hypoxia (47), it can also reduce VEGF levels (80), and suppress key glycolytic enzymes and reduce mitochondrial membrane potential (81) (82).

Our data indicate that HIF-1 activation alone is insufficient to drive autophagic flux in the MFs. Although DMOG markedly increased HIF1α levels, it did not significantly alter LC3-II or p62 protein levels. Similarly, while LW6 reduced the expression of autophagy-related genes, DMOG failed to increase their expression, except for *BNIP3*. BNIP3, a HIF-regulated pro-apoptotic BH3-only member of the BCL2 protein family, which plays a crucial role in autophagy by facilitating mitophagy and promoting autophagosome formation through its interaction with LC3 (83). These findings suggest that additional factors, such as hypoxia, may be required to fully stimulate autophagy under pharmacological HIF activation, and genetic perturbation studies could further validate the findings related to HIF1α. Nonetheless, our results reveal a complex regulatory axis in which lactate stabilizes HIF1α in MFs, promoting an autophagic state.

A limitation of this study is that the downstream mediators by which autophagy-competent MFs promote cyst epithelial cell proliferation were not defined. While our data demonstrate that autophagy in MFs is required for their pro-proliferative paracrine effects, the specific factors involved remain unknown. These effects may be mediated by changes in extracellular matrix or soluble signals derived from MFs. Identifying the autophagy-dependent mechanisms that drive this epithelial–mesenchymal crosstalk could be a focus of future studies.

## Conclusion

Our findings identify autophagy in MFs as a critical driver of cyst growth and kidney fibrosis in ADPKD. Targeting autophagy in MFs may represent a promising therapeutic strategy to limit cyst expansion and fibrotic remodeling.

## Supporting information

Unedited western blot images

## List of abbreviations

ADPKD: Autosomal dominant polycystic kidney disease
Atg5: Autophagy related gene-5
HIF1α: Hypoxia-inducible factor-1α
WT: wild type
ECM: Extracellular matrix
MFs: Myofibroblasts
ADPKD-MFs: Human ADPKD kidney Myofibroblasts
ADPKD-ECs: Human ADPKD kidney cyst epithelial cells
NHK: Normal human kidney
NHK-ECs: Normal human kidney epithelial cells
αSMA: α-smooth muscle actin
CKD: Chronic kidney disease
LC3: microtubule-asociated protein light chain 3
RC/RC: Pkd1^RC/RC^
CQ: Chloroquine
CM: conditioned media
Fn1: Fibronectin
3-HBA: 3-Hydroxybutyric acid
α-KGA: α-ketoglutaric acid
PHDs: prolyl hydroxylases
DMOG: dimethyloxalylglycine

## Ethics declarations

### Ethics approval

All animal experiments received approval from the Institutional Animal Care and Use Committee of the University of Kansas.

### Consent for publication

Not applicable.

## Acknowledgments

We thank Dr. Peter Harris and Dr. Katharina Hopp for providing *Pkd1*^RC/RC^ mice. *Atg5*^f/f^ mice were originally generated by Dr. Noboru Mizushima, University of Tokyo and the RIKEN BRC, Japan, and supplied Dr. Ning Wang. Human kidney tissues and primary cultures of NHK and ADPKD epithelial cells were obtained from the NIDDK PKD-Research Resource Consortium and the Kansas PKD Research and Translational Core Center (U54 DK126126). We acknowledge the University of Kansas Medical Center’s Imaging core and histology core and the flow cytometry core Laboratory, which is sponsored, in part, by the NIH/NIGMS COBRE grant P30 GM103326 and the NIH/NCI Cancer Center grant P30 CA168524. We acknowledge Penn Metabolomics Core (RRID:SCR_022381) supported by the Penn Cardiovascular Institute and, in part, by NCI P30 CA016520 and NIH P30DK050306.

## Author contributions

RR conceptualized and designed studies, analyzed results, and wrote the paper. AJ performed experiments, analyzed results and wrote parts of the paper. VR, MMS, HY, IC, VPT performed some studies and analyzed results. DPW, WXD and VHH conceptualized some studies and reviewed the paper. All authors read, edited and approved the paper.

## Data sharing statements

No proteomics, transcriptomics or GWAS data was generated in this paper. All *in vivo* studies using mouse models were performed and tissues analyzed at the University of Kansas Medical Center. All data that was generated are provided in the figures in the main text or in Supplementary Material.

All requests for data will be processed based on institutional policies for noncommercial research purposes. Data sharing could require a data transfer agreement as determined by the University of Kansas Medical Center’s legal department.

## REFERENCES

1. Pei Y, Watnick T. Diagnosis and screening of autosomal dominant polycystic kidney disease. Adv Chronic Kidney Dis. 2010;17(2):140–52.

2. Chapman AB, Devuyst O, Eckardt KU, Gansevoort RT, Harris T, Horie S, et al. Autosomal-dominant polycystic kidney disease (ADPKD): executive summary from a Kidney Disease: Improving Global Outcomes (KDIGO) Controversies Conference. Kidney Int. 2015.

3. Fragiadaki M, Macleod FM, Ong ACM. The Controversial Role of Fibrosis in Autosomal Dominant Polycystic Kidney Disease. Int J Mol Sci. 2020;21(23).

4. Song CJ, Zimmerman KA, Henke SJ, Yoder BK. Inflammation and Fibrosis in Polycystic Kidney Disease. Results Probl Cell Differ. 2017;60:323–44.

5. Xue C, Mei CL. Polycystic Kidney Disease and Renal Fibrosis. Adv Exp Med Biol. 2019;1165:81–100.

6. Duffield JS. Cellular and molecular mechanisms in kidney fibrosis. J Clin Invest. 2014;124(6):2299–306.

7. Eddy AA. Molecular basis of renal fibrosis. Pediatr Nephrol. 2000;15(3-4):290–301.

8. Farris AB, Colvin RB. Renal interstitial fibrosis: mechanisms and evaluation. Curr Opin Nephrol Hypertens. 2012;21(3):289–300.

9. Grgic I, Duffield JS, Humphreys BD. The origin of interstitial myofibroblasts in chronic kidney disease. Pediatr Nephrol. 2012;27(2):183–93.

10. LeBleu VS, Taduri G, O’Connell J, Teng Y, Cooke VG, Woda C, et al. Origin and function of myofibroblasts in kidney fibrosis. Nat Med. 2013;19(8):1047–53.

11. Otranto M, Sarrazy V, Bonte F, Hinz B, Gabbiani G, Desmouliere A. The role of the myofibroblast in tumor stroma remodeling. Cell Adh Migr. 2012;6(3):203–19.

12. Dwivedi N, Jamadar A, Mathew S, Fields TA, Rao R. Myofibroblast depletion reduces kidney cyst growth and fibrosis in autosomal dominant polycystic kidney disease. Kidney Int. 2023;103(1):144–55.

13. Dwivedi N, Tao S, Jamadar A, Sinha S, Howard C, Wallace DP, et al. Epithelial Vasopressin Type-2 Receptors Regulate Myofibroblasts by a YAP-CCN2-Dependent Mechanism in Polycystic Kidney Disease. J Am Soc Nephrol. 2020;31(8):1697–710.

14. Jamadar A, Suma SM, Mathew S, Fields TA, Wallace DP, Calvet JP, et al. The tyrosine-kinase inhibitor Nintedanib ameliorates autosomal-dominant polycystic kidney disease. Cell Death Dis. 2021;12(10):947.

15. Remadevi V, Jamadar A, Varghese MM, Gunewardena S, Wallace D, Rao R. Pirfenidone treatment attenuates fibrosis in autosomal dominant polycystic kidney disease. bioRxiv. 2025:2025.08.25.672225.

16. Kuma A, Hatano M, Matsui M, Yamamoto A, Nakaya H, Yoshimori T, et al. The role of autophagy during the early neonatal starvation period. Nature. 2004;432(7020):1032–6.

17. Cuervo AM, Macian F. Autophagy, nutrition and immunology. Mol Aspects Med. 2012;33(1):2–13.

18. Janku F, McConkey DJ, Hong DS, Kurzrock R. Autophagy as a target for anticancer therapy. Nat Rev Clin Oncol. 2011;8(9):528–39.

19. Neikirk K, Vue Z, Katti P, Rodriguez BI, Omer S, Shao J, et al. Systematic Transmission Electron Microscopy-Based Identification and 3D Reconstruction of Cellular Degradation Machinery. Adv Biol (Weinh). 2023;7(6):e2200221.

20. Chiaravalli M, Rowe I, Mannella V, Quilici G, Canu T, Bianchi V, et al. 2-Deoxy-d-Glucose Ameliorates PKD Progression. J Am Soc Nephrol. 2016;27(7):1958–69.

21. Ravichandran K, Edelstein CL. Polycystic kidney disease: a case of suppressed autophagy? Semin Nephrol. 2014;34(1):27–33.

22. Zhu P, Sieben CJ, Xu X, Harris PC, Lin X. Autophagy activators suppress cystogenesis in an autosomal dominant polycystic kidney disease model. Hum Mol Genet. 2017;26(1):158–72.

23. Lee EJ, Ko JY, Oh S, Jun J, Mun H, Lim CJ, et al. Autophagy induction promotes renal cyst growth in polycystic kidney disease. EBioMedicine. 2020;60:102986.

24. Martinez-Outschoorn UE, Whitaker-Menezes D, Pavlides S, Chiavarina B, Bonuccelli G, Casey T, et al. The autophagic tumor stroma model of cancer or “battery-operated tumor growth”: A simple solution to the autophagy paradox. Cell Cycle. 2010;9(21):4297–306.

25. Grimwood L, Masterson R. Propagation and culture of renal fibroblasts. Methods Mol Biol. 2009;466:25–37.

26. Hahn VS, Petucci C, Kim MS, Bedi KC, Jr., Wang H, Mishra S, et al. Myocardial Metabolomics of Human Heart Failure With Preserved Ejection Fraction. Circulation. 2023;147(15):1147–61.

27. Hopp K, Ward CJ, Hommerding CJ, Nasr SH, Tuan HF, Gainullin VG, et al. Functional polycystin-1 dosage governs autosomal dominant polycystic kidney disease severity. J Clin Invest. 2012;122(11):4257–73.

28. Cuervo H, Pereira B, Nadeem T, Lin M, Lee F, Kitajewski J, et al. PDGFRbeta-P2A-CreER(T2) mice: a genetic tool to target pericytes in angiogenesis. Angiogenesis. 2017;20(4):655–62.

29. Percie du Sert N, Hurst V, Ahluwalia A, Alam S, Avey MT, Baker M, et al. The ARRIVE guidelines 2.0: Updated guidelines for reporting animal research. PLoS Biol. 2020;18(7):e3000410.

30. Jamadar A, Dwivedi N, Mathew S, Calvet JP, Thomas SM, Rao R. Vasopressin Receptor Type-2 Mediated Signaling in Renal Cell Carcinoma Stimulates Stromal Fibroblast Activation. International Journal of Molecular Sciences. 2022;23(14):7601.

31. Sinha S, Dwivedi N, Tao S, Jamadar A, Kakade VR, Neil MO, et al. Targeting the vasopressin type-2 receptor for renal cell carcinoma therapy. Oncogene. 2020;39(6):1231–45.

32. Abrahamson DR, St John PL. Loss of laminin epitopes during glomerular basement membrane assembly in developing mouse kidneys. J Histochem Cytochem. 1992;40(12):1943–53.

33. Jamadar A, Ward CJ, Remadevi V, Varghese MM, Pabla NS, Gumz ML, et al. Circadian Clock Disruption and Growth of Kidney Cysts in Autosomal Dominant Polycystic Kidney Disease. J Am Soc Nephrol. 2024.

34. Norregaard R, Tao S, Nilsson L, Woodgett JR, Kakade V, Yu AS, et al. Glycogen synthase kinase 3alpha regulates urine concentrating mechanism in mice. Am J Physiol Renal Physiol. 2015;308(6):F650–60.

35. Singh SP, Tao S, Fields TA, Webb S, Harris RC, Rao R. Glycogen synthase kinase-3 inhibition attenuates fibroblast activation and development of fibrosis following renal ischemia-reperfusion in mice. Dis Model Mech. 2015;8(8):931–40.

36. Busselman BW, Ratnayake I, Terasaki MR, Thakkar VP, Ilyas A, Otterpohl KL, et al. Actin cytoskeleton and associated myosin motors within the renal epithelium. Am J Physiol Renal Physiol. 2024;327(4):F553–F65.

37. Kabeya Y, Mizushima N, Ueno T, Yamamoto A, Kirisako T, Noda T, et al. LC3, a mammalian homologue of yeast Apg8p, is localized in autophagosome membranes after processing. EMBO J. 2000;19(21):5720–8.

38. Mizushima N, Yoshimori T. How to interpret LC3 immunoblotting. Autophagy. 2007;3(6):542–5.

39. Livingston MJ, Ding HF, Huang S, Hill JA, Yin XM, Dong Z. Persistent activation of autophagy in kidney tubular cells promotes renal interstitial fibrosis during unilateral ureteral obstruction. Autophagy. 2016;12(6):976–98.

40. Buhl EM, Djudjaj S, Klinkhammer BM, Ermert K, Puelles VG, Lindenmeyer MT, et al. Dysregulated mesenchymal PDGFR-beta drives kidney fibrosis. EMBO Mol Med. 2020;12(3):e11021.

41. Arroyo J, Escobar-Zarate D, Wells HH, Constans MM, Thao K, Smith JM, et al. The genetic background significantly impacts the severity of kidney cystic disease in the Pkd1(RC/RC) mouse model of autosomal dominant polycystic kidney disease. Kidney Int. 2021;99(6):1392–407.

42. Ni HM, Boggess N, McGill MR, Lebofsky M, Borude P, Apte U, et al. Liver-specific loss of Atg5 causes persistent activation of Nrf2 and protects against acetaminophen-induced liver injury. Toxicol Sci. 2012;127(2):438–50.

43. Kraus A, Peters DJM, Klanke B, Weidemann A, Willam C, Schley G, et al. HIF-1alpha promotes cyst progression in a mouse model of autosomal dominant polycystic kidney disease. Kidney Int. 2018;94(5):887–99.

44. Deng J, Huang Q, Wang Y, Shen P, Guan F, Li J, et al. Hypoxia-inducible factor-1alpha regulates autophagy to activate hepatic stellate cells. Biochem Biophys Res Commun. 2014;454(2):328–34.

45. Deng M, Zhang W, Yuan L, Tan J, Chen Z. HIF-1a regulates hypoxia-induced autophagy via translocation of ANKRD37 in colon cancer. Exp Cell Res. 2020;395(1):112175.

46. De Saedeleer CJ, Copetti T, Porporato PE, Verrax J, Feron O, Sonveaux P. Lactate activates HIF-1 in oxidative but not in Warburg-phenotype human tumor cells. PLoS One. 2012;7(10):e46571.

47. Lee K, Kang JE, Park SK, Jin Y, Chung KS, Kim HM, et al. LW6, a novel HIF-1 inhibitor, promotes proteasomal degradation of HIF-1alpha via upregulation of VHL in a colon cancer cell line. Biochem Pharmacol. 2010;80(7):982–9.

48. Qian H, Chao X, Williams J, Fulte S, Li T, Yang L, et al. Autophagy in liver diseases: A review. Mol Aspects Med. 2021;82:100973.

49. Tang C, Livingston MJ, Liu Z, Dong Z. Autophagy in kidney homeostasis and disease. Nat Rev Nephrol. 2020;16(9):489–508.

50. Atwood DJ, Brown CN, Holditch SJ, Pokhrel D, Thorburn A, Hopp K, et al. The effect of trehalose on autophagy-related proteins and cyst growth in a hypomorphic Pkd1 mouse model of autosomal dominant polycystic kidney disease. Cell Signal. 2020;75:109760.

51. Chou LF, Cheng YL, Hsieh CY, Lin CY, Yang HY, Chen YC, et al. Effect of Trehalose Supplementation on Autophagy and Cystogenesis in a Mouse Model of Polycystic Kidney Disease. Nutrients. 2018;11(1).

52. Ghavami S, Cunnington RH, Gupta S, Yeganeh B, Filomeno KL, Freed DH, et al. Autophagy is a regulator of TGF-beta1-induced fibrogenesis in primary human atrial myofibroblasts. Cell Death Dis. 2015;6(3):e1696.

53. Gupta SS, Zeglinski MR, Rattan SG, Landry NM, Ghavami S, Wigle JT, et al. Inhibition of autophagy inhibits the conversion of cardiac fibroblasts to cardiac myofibroblasts. Oncotarget. 2016;7(48):78516–31.

54. Rudnick JA, Monkkonen T, Mar FA, Barnes JM, Starobinets H, Goldsmith J, et al. Autophagy in stromal fibroblasts promotes tumor desmoplasia and mammary tumorigenesis. Genes Dev. 2021;35(13-14):963–75.

55. Tan ML, Parkinson EK, Yap LF, Paterson IC. Autophagy is deregulated in cancer-associated fibroblasts from oral cancer and is stimulated during the induction of fibroblast senescence by TGF-beta1. Sci Rep. 2021;11(1):584.

56. Zhang X, Lao M, Yang H, Sun K, Dong Y, He L, et al. Targeting cancer-associated fibroblast autophagy renders pancreatic cancer eradicable with immunochemotherapy by inhibiting adaptive immune resistance. Autophagy. 2024;20(6):1314–34.

57. Formica C, Peters DJM. Molecular pathways involved in injury-repair and ADPKD progression. Cell Signal. 2020;72:109648.

58. Nigro EA, Boletta A. Role of the polycystins as mechanosensors of extracellular stiffness. Am J Physiol Renal Physiol. 2021;320(5):F693–F705.

59. Zhang Y, Reif G, Wallace DP. Extracellular matrix, integrins, and focal adhesion signaling in polycystic kidney disease. Cell Signal. 2020;72:109646.

60. Karihaloo A, Koraishy F, Huen SC, Lee Y, Merrick D, Caplan MJ, et al. Macrophages promote cyst growth in polycystic kidney disease. J Am Soc Nephrol. 2011;22(10):1809–14.

61. Kleczko EK, Marsh KH, Tyler LC, Furgeson SB, Bullock BL, Altmann CJ, et al. CD8(+) T cells modulate autosomal dominant polycystic kidney disease progression. Kidney Int. 2018;94(6):1127–40.

62. Swenson-Fields KI, Vivian CJ, Salah SM, Peda JD, Davis BM, van Rooijen N, et al. Macrophages promote polycystic kidney disease progression. Kidney Int. 2013;83(5):855–64.

63. Zimmerman KA, Gonzalez NM, Chumley P, Chacana T, Harrington LE, Yoder BK, et al. Urinary T cells correlate with rate of renal function loss in autosomal dominant polycystic kidney disease. Physiol Rep. 2019;7(1):e13951.

64. Zimmerman KA, Song CJ, Li Z, Lever JM, Crossman DK, Rains A, et al. Tissue-Resident Macrophages Promote Renal Cystic Disease. J Am Soc Nephrol. 2019;30(10):1841–56.

65. Hsieh CL, Jerman SJ, Sun Z. Non-cell-autonomous activation of hedgehog signaling contributes to disease progression in a mouse model of renal cystic ciliopathy. Hum Mol Genet. 2022;31(24):4228–40.

66. Lichner Z, Ding M, Khare T, Dan Q, Benitez R, Praszner M, et al. Myocardin-Related Transcription Factor Mediates Epithelial Fibrogenesis in Polycystic Kidney Disease. Cells. 2024;13(11).

67. Podrini C, Rowe I, Pagliarini R, Costa ASH, Chiaravalli M, Di Meo I, et al. Dissection of metabolic reprogramming in polycystic kidney disease reveals coordinated rewiring of bioenergetic pathways. Commun Biol. 2018;1:194.

68. Rowe I, Chiaravalli M, Mannella V, Ulisse V, Quilici G, Pema M, et al. Defective glucose metabolism in polycystic kidney disease identifies a new therapeutic strategy. Nat Med. 2013;19(4):488–93.

69. Jia M, Yue X, Sun W, Zhou Q, Chang C, Gong W, et al. ULK1-mediated metabolic reprogramming regulates Vps34 lipid kinase activity by its lactylation. Sci Adv. 2023;9(22):eadg4993.

70. Su BC, Hsu PL, Mo FE. CCN1 triggers adaptive autophagy in cardiomyocytes to curb its apoptotic activities. J Cell Commun Signal. 2020;14(1):93–100.

71. Ni Y, Zhang H, Li Z, Li Z. Connective tissue growth factor (CCN2) inhibits TNF-alpha-induced apoptosis by enhancing autophagy through the Akt and Erk pathways in osteoblasts. Pharmazie. 2020;75(5):213–7.

72. Belibi F, Zafar I, Ravichandran K, Segvic AB, Jani A, Ljubanovic DG, et al. Hypoxia-inducible factor-1alpha (HIF-1alpha) and autophagy in polycystic kidney disease (PKD). Am J Physiol Renal Physiol. 2011;300(5):F1235–43.

73. Buchholz B, Schley G, Faria D, Kroening S, Willam C, Schreiber R, et al. Hypoxia-inducible factor-1alpha causes renal cyst expansion through calcium-activated chloride secretion. J Am Soc Nephrol. 2014;25(3):465–74.

74. Theodorakopoulou M, Raptis V, Loutradis C, Sarafidis P. Hypoxia and Endothelial Dysfunction in Autosomal-Dominant Polycystic Kidney Disease. Semin Nephrol. 2019;39(6):599–612.

75. Eckardt KU, Mollmann M, Neumann R, Brunkhorst R, Burger HU, Lonnemann G, et al. Erythropoietin in polycystic kidneys. J Clin Invest. 1989;84(4):1160–6.

76. Grantham JJ. Clinical practice. Autosomal dominant polycystic kidney disease. N Engl J Med. 2008;359(14):1477–85.

77. Ibrahim S. Increased apoptosis and proliferative capacity are early events in cyst formation in autosomal-dominant, polycystic kidney disease. ScientificWorldJournal. 2007;7:1757–67.

78. Norman J. Fibrosis and progression of autosomal dominant polycystic kidney disease (ADPKD). Biochim Biophys Acta. 2011;1812(10):1327–36.

79. Schodel J, Ratcliffe PJ. Mechanisms of hypoxia signalling: new implications for nephrology. Nat Rev Nephrol. 2019;15(10):641–59.

80. Xu H, Chen Y, Li Z, Zhang H, Liu J, Han J. The hypoxia-inducible factor 1 inhibitor LW6 mediates the HIF-1alpha/PD-L1 axis and suppresses tumor growth of hepatocellular carcinoma in vitro and in vivo. Eur J Pharmacol. 2022;930:175154.

81. Eleftheriadis T, Pissas G, Antoniadi G, Liakopoulos V, Stefanidis I. Malate dehydrogenase-2 inhibitor LW6 promotes metabolic adaptations and reduces proliferation and apoptosis in activated human T-cells. Exp Ther Med. 2015;10(5):1959–66.

82. Sato M, Hirose K, Kashiwakura I, Aoki M, Kawaguchi H, Hatayama Y, et al. LW6, a hypoxia-inducible factor 1 inhibitor, selectively induces apoptosis in hypoxic cells through depolarization of mitochondria in A549 human lung cancer cells. Mol Med Rep. 2015;12(3):3462–8.

83. Zhang J, Ney PA. Role of BNIP3 and NIX in cell death, autophagy, and mitophagy. Cell Death Differ. 2009;16(7):939–46.

