## Supplementary File for "Myofibroblast- specific autophagy drives cyst growth in autosomal dominant polycystic kidney disease"

**SUPPLEMENTARY MATERIAL:**

**Table of contents:**

**Supplementary video 1 (Attached as a separate avi file)**

**Supplementary Figure 1**

**Supplementary Figure 2**

**Supplementary Figure 3**

**Supplementary Figure 4**

**Supplementary Figure 5**

**Supplementary Figure 6**

**Supplementary Figure 7**

**Supplementary Figure 8**

**Supplementary Table 1**

**Supplementary Figure Legends**

**Supplementary methods**

**Uncut Western blot images (Attached as separate pdf file)**

**Supplementary Excel File-1(Attached as separate excel file)**

Supplementary Fig 1

(A)

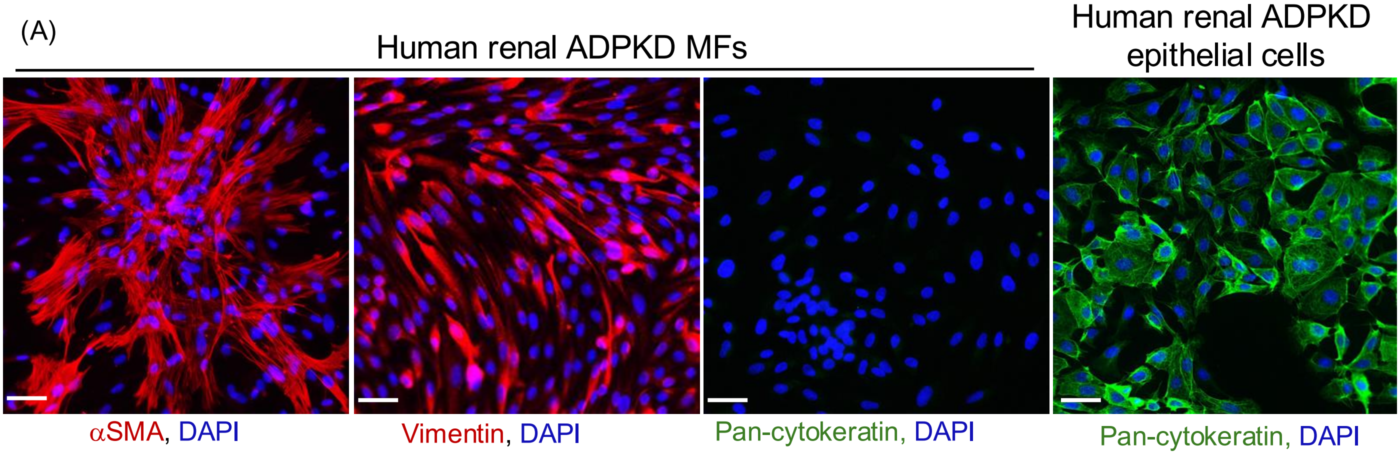


(B)

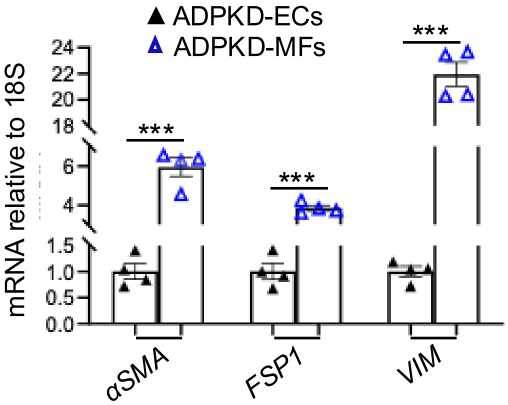


(C)

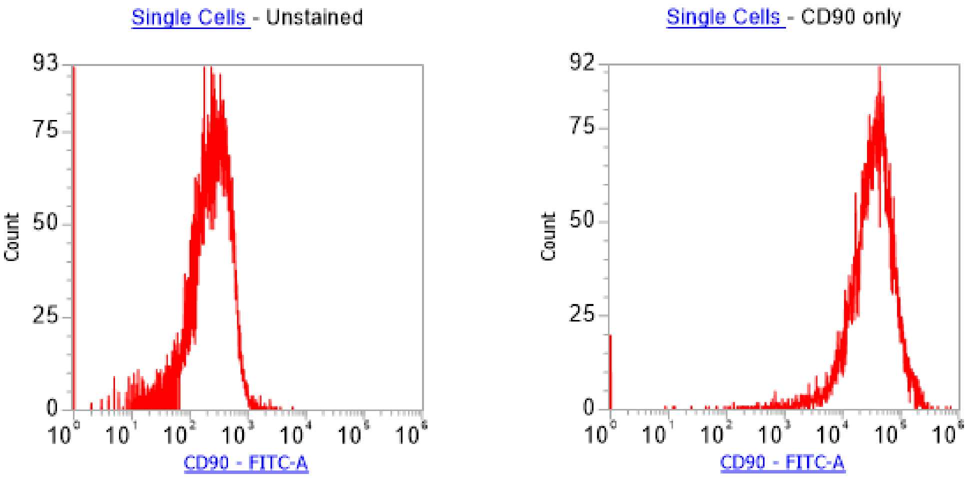

Supplementary Fig 3

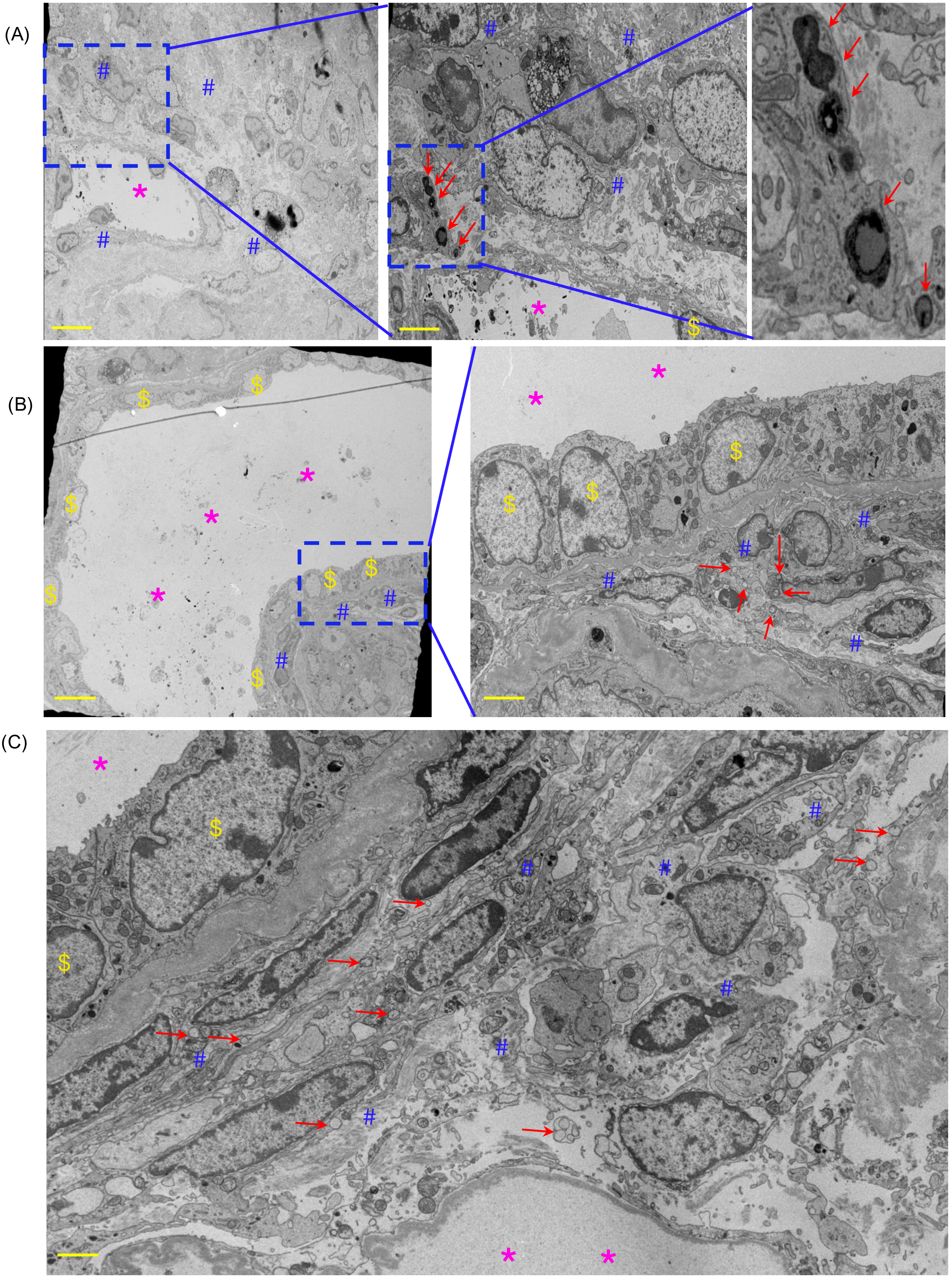

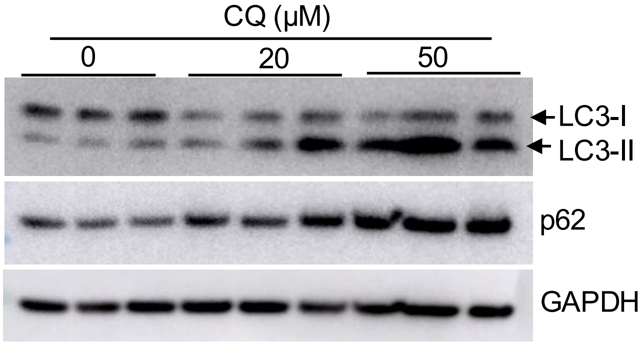
CQ (μM)
0
20
50
LC3-I
LC3-II
p62
GAPDH

(A)

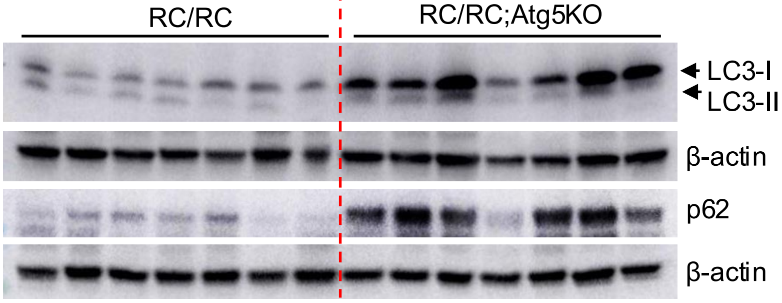
RC/RC
RC/RC;Atg5KO
LC3-I
LC3-II
β-actin
p62
β-actin

(B)

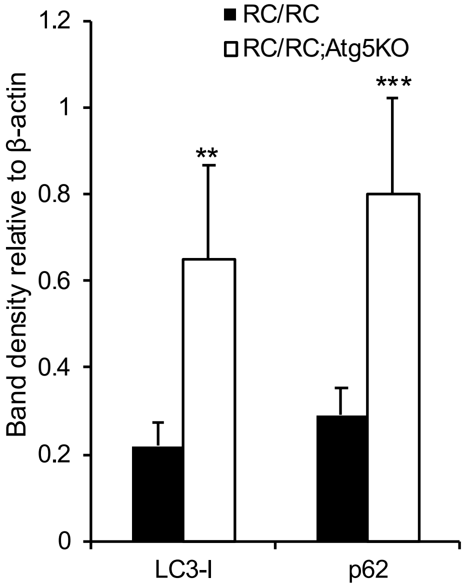
RC/RC
RC/RC;Atg5KO
Band density relative to β-actin
1.2
1
0.8
0.6
0.4
0.2
0
**
***
LC3-I
p62

(C)

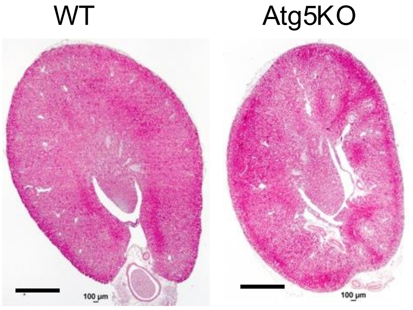
WT
Atg5KO

(A)

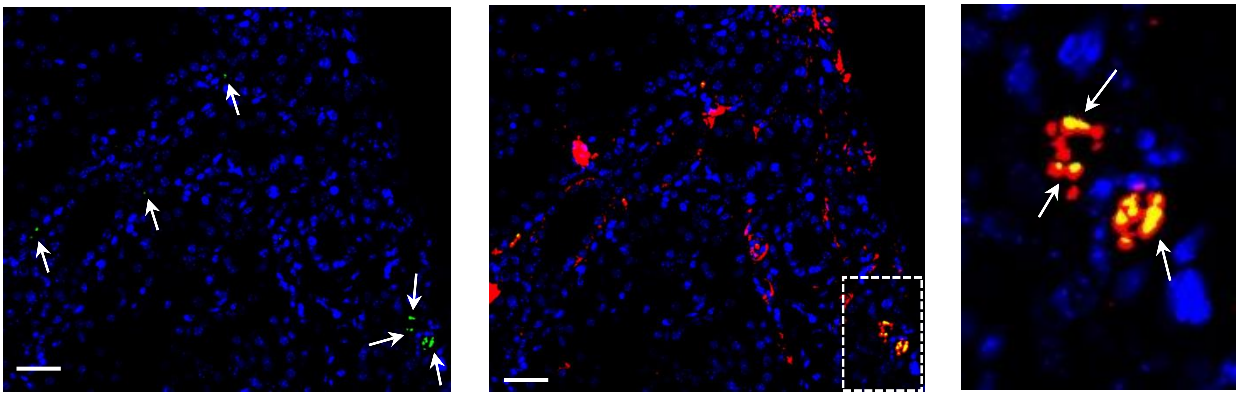

LC3, αSMA, DAPI

(B)

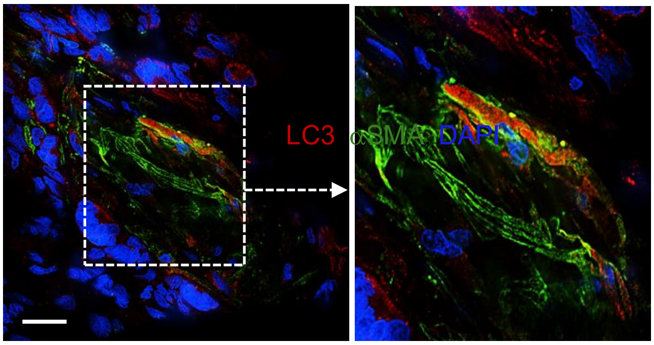

LC3, αSMA, DAPI

(C)

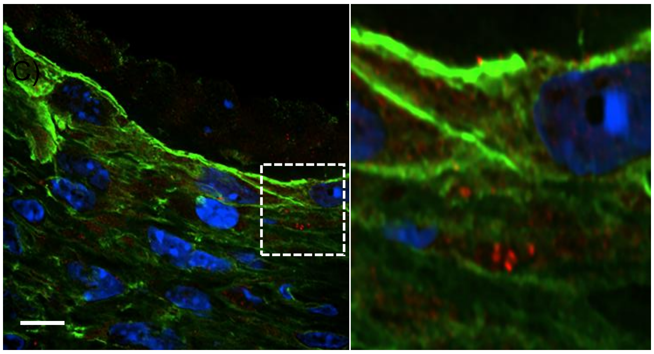

LC3, αSMA, DAPI

Supplementary Fig 6

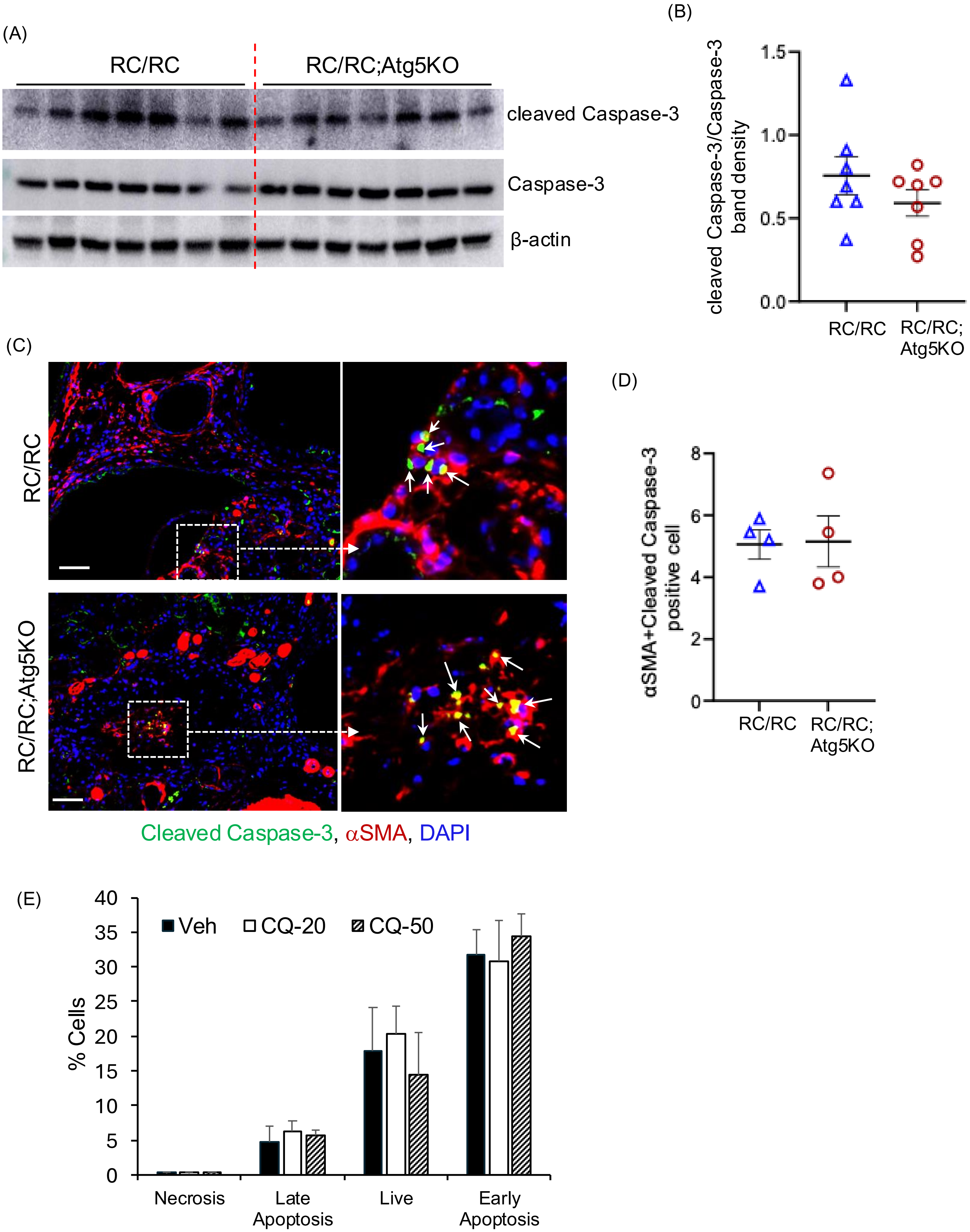

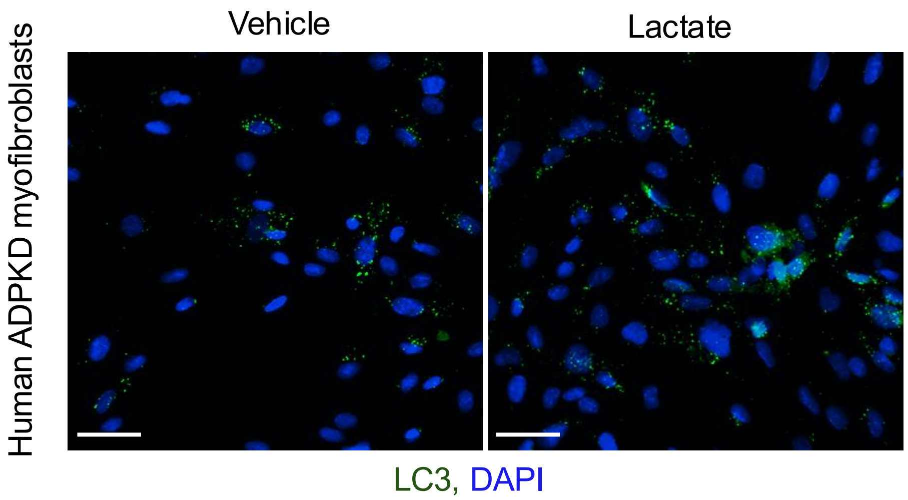
Vehicle
Lactate
Human ADPKD myofibroblasts
LC3, DAPI

(A)

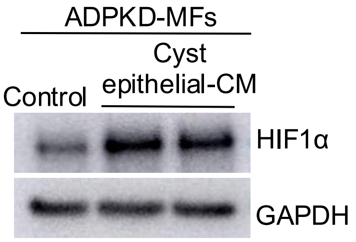
ADPKD-MFs
Control
Cyst epithelial-CM
HIF1α
GAPDH

(B)

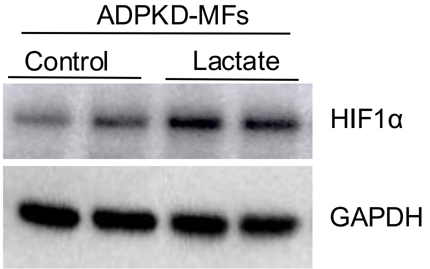
ADPKD-MFs
Control
Lactate
HIF1α
GAPDH

(C)

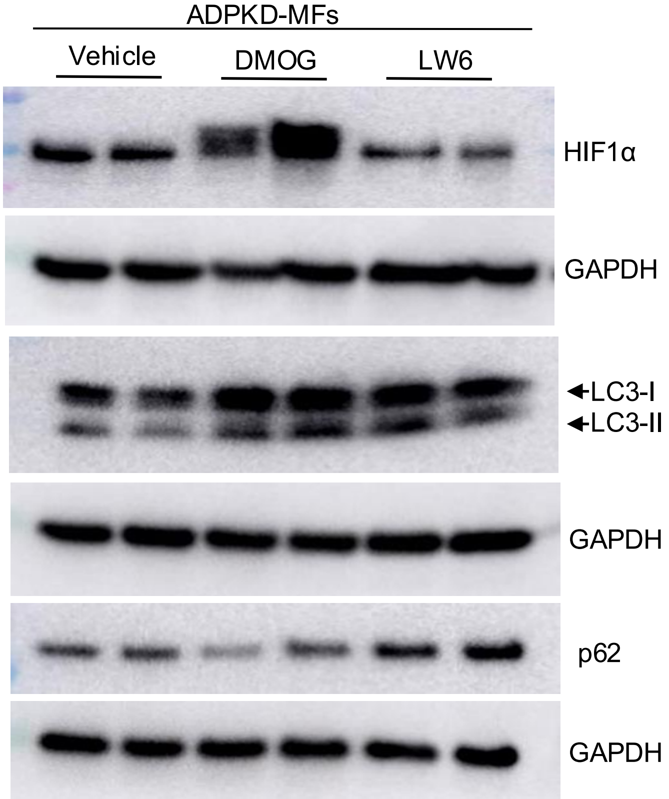
ADPKD-MFs
Vehicle
DMOG
LW6
HIF1α
GAPDH
LC3-I
LC3-II
GAPDH
p62
GAPDH

**Supplementary Figure legend:**

**Supplementary Figure 1:** ADPKD-MFs immunostained for (A) αSMA (red), vimentin (red), pan-cytokeratin (green) and DAPI (blue) and ADPKD-ECs immunostained for pan-cytokeratin (green) and DAPI (blue) (Scale bar = 200μm) (D) ADPKD-ECs immunostained pan-cytokeratin (green) and DAPI (blue) (Scale bar = 200μm). (B) QRTPCR of ADPKD MFs and ADPKD ECs. (C) Histogram representation of flow cytometry analysis for CD90 (Thy1) expression of ADPKD MFs. ***$P<0.001$by T-test for B.

**Supplementary Figure 2:** (A) Immunostaining for αSMA (red, denotes MFs), LC3 (green, denotes autophagosome) and DAPI (blue, denotes nuclei) in kidney tissue sections of 4 months old RC/RC mice. The arrows points to LC3 puncta. (Scale bar = 200 μm). High resolution microscopy images of (B) human ADPKD kidney, and (C) RC/RC mouse kidney tissue sections immunostained for αSMA (green, denotes MFs), LC3 (red, denotes autophagosome) and DAPI (blue) (Scale bar = 5 μm).

**Supplementary Figure 3:** (A), (B), (C) Transmission electron microscopy images of RC/RC mouse kidney sections of 6.5 months of age, red arrows indicates autophagic organelles, * denotes cyst lumen area, # denotes interstitium and $ denotes cyst lining epithelial cells (Scale 10 μm and 2 μm for A, B and 2 μm for C).

**Supplementary Figure 4:** Immunoblot of ADPKD-MFs treated with 20μM and 50μM chloroquine (CQ) for 24 hours.

**Supplementary Figure 5:** (A) Immunoblot on mouse kidney tissue lysate and its (B) Quantitation of band density of Supplementary Fig 4B. (C) H&E-stained kidney tissue sections (Scale bar = 1mm). **P<0.01, ***P<0.001by T-test for B.

**Supplementary Figure 6:** (A) Immunoblot on kidney tissue lysate and (B) Quantitation of band density for cleaved Caspase-3/Caspase-3 for supplementary Fig 4A. (C) Immunostaining of cleaved Caspase-3 (green), αSMA (red) and DAPI (blue) in RC/RC and RC/RC;Atg5 in mice kidney sections (Scale bar = 200μm). (D) Average number of cleaved Caspase-3 in αSMA expressing cells per kidney section of individual mice. (E) ADPKD-MFs treated with CQ for 24h. Annexin/PI were analyzed by flow cytometry.

**Supplementary Figure 7:** Immunostaining for LC3 (green) and DAPI (blue), (Scale bar = 200μm) in ADPKD-MFs exposed to lactate (20 mM for 24 hours, Scale bar = 200μm).

**Supplementary Figure 8:** Immunoblot for ADPKD-MFs treated with (A) ADPKD-ECs CM (B) lactate (20 mM) and (C) DMOG (1mM) or LW6 (20 μM) for 24 hours.

**Supplemental methods:**

**Collection of conditioned media:**

Monolayers of human primary culture ADPKD kidney epithelial cells or myofibroblasts were grown in 100 mm petri dishes. To collect cell culture conditioned media, confluent monolayers of cells were incubated in serum free DMEM/F12 with 1% pen/strip and media was collected after 24 hours. The conditioned media was centrifuged at 2000 rpm for 5 minutes to remove cell debris. Serum free media (DMEM/F12 with 1% pen/strip) not exposed to any cells was used as control.

***In vitro* BrdU incorporation:**

20,000 human primary culture epithelial cells were seeded on glass coverslips and grown in DMEM:F12 media containing 5% FBS and 1% pen/strep. 40-50% confluent cells were incubated with FBS-free media for 16h followed by incubation with cell culture conditioned media with and without 20µM chloroquine for 24h. Cells were incubated with BrdU (#10280879001, MilliporeSigma, Burlington, MA) (3 µg/mL) for the last 3 h of incubation (1). Cells were fixed and immunostained for BrdU (#5292S, Cell Signaling Technology, Danvers, MA) ad DAPI. Images were taken using Nikon 80i microscope and quantified using Image J software.

**Detection of apoptosis/necrosis using Annexin V-FITC/PI staining:**

Annexin V-FITC/PI staining was performed using FITC Annexin V apoptosis detection kit (#130-092-052, Miltenyi biotec B.V & Co.KG, Bergisch Gladbach, Germany). 2 X $10^5$ human primary culture fibroblasts were seeded in a 60 mm plates. The cells were serum starved overnight followed by treatment with vehicle or chloroquine (20 and 50 $\mu$M) in 0.2 % FBS containing media for 24 hours. At the end of the study, the cells were washed in ice-cold PBS, resuspended in calcium containing binding buffer (10 mM

HEPES, 140mM NaCl, 5mM CaCl2; pH 7.4) and stained for 15min, with 5µl Annexin V-FITC and 5µl PI at 1µg/ml. The stained cells were then analyzed for FITC (excitation 488nm and emission 530nm) and PI (610nm) fluorescence using AttuneNxT flow cytometer. CellQuestPro® software (Becton, Dickinson, Heidelberg, Germany) was used to determine the percent viable (Annexin V-PI-), early apoptotic (Annexin V+PI-), late apoptotic/necrotic (Annexin V+PI+) and necrotic cells (Annexin V-PI+).

**Flow Cytometric analysis if ADPKD myofibroblasts for CD90+ (thy1) positive cells:**

ADPKD MFs were trypsinized and centrifuged for 2 min at 2000 rpm. The supernatant was discarded by aspiration, and the cells were washed twice with flow cytometer buffer (phosphate-buffered saline (PBS), 2 % FBS) and incubated for 30 min in dark with containing CD90-FITC (#328108, BioLegend, San Diego, California, US). The cells were washed twice with flow cytometer buffer and stained cells were then analyzed for FITC (excitation 488nm and emission 530nm) fluorescence using AttuneNxT flow cytometer.

**Western blot:**

Mouse kidney tissues were homogenized in SDS Laemmli buffer and loaded onto SDS polyacrylamide agarose electrophoresis gels as described previously (2, 3). Primary antibodies used were, LC3 I/II (#12741) and p62 (#5114) from Cell signaling (Danvers, MA, USA); HIf1α (#ab179483) and αSMA (#ab5694) from Abcam (Cambridge, MA) and GAPDH (SC-32233) from Santa Cruz Biotechnology, Inc. (Dallas, TX, USA). The secondary antibodies for anti-rabbit (#P0448) and anti-mouse (#P0447) were purchased from Dako (Santa Clara, CA, USA). ECL reagent was purchased from Amersham (GE Healthcare, Buckinghamshire, UK).

**Immunohistochemistry/ immunofluorescence (IHC/ IF) staining:**

For IHC, the mouse kidney tissues were fixed in 4% paraformaldehyde (#50-980-487, Electron Microscopy Sciences, Hatfield, PA, USA) and embedded in paraffin as described before (4). For IHC, the primary antibodies were, LC3 (#12741) and Ki-67 (#12202) from Cell Signaling Technology, Danvers, MA. Tissue sections were incubated with streptavidin HRP conjugate secondary antibody (Invitrogen, Carlsbad, CA, USA), followed by DAB (Vector Laboratories, Burlingame, CA, USA). Following counterstaining with Harris Hematoxylin stain, tissues were dehydrated, and mounted using Permount (Fisher Scientific, Waltham, MA, USA).

For IF, LC3 (#12741) from Cell signaling (Danvers, MA, USA) and HIf1$\alpha$ (#ab179483), $\alpha$SMA (#ab5694) from Abcam (Cambridge, MA), Vimentin (SC7557) from Santa Cruz Biotechnology, Inc. (Dallas, TX, USA), and Pan-cytokeratin (F0397) from (MilliporeSigma, St. Louis, MO) primary antibodies were used. Secondary antibodies used were goat anti-Rabbit IgG fluor and goat anti-mouse IgG Texas red (Invitrogen, Carlsbad, CA, USA). After incubation with secondary antibodies, tissue sections were washed and stained with DAPI, and mounted with Flour-G (Invitrogen, Carlsbad, CA, USA). Images were taken using a Nikon 90i upright microscope (Tokyo, Japan).

**Super resolution microscopy- vibratome sectioning and immunostaining protocol:**

The mouse kidneys were fixed in 4 % PFA (1:1 sodium cacodylate buffer, 0.2M, pH 7.4 #11650 and 8% PFA #157-8-100, Electron Microscopy Sciences, Hatfield, PA, USA) for

4 days followed by replacing with 70% ethanol until further processing. The kidneys were cut into half sagitally using a sharp blade. One of the halves was embedded in a plastic embedding mold with 4.0% w/v agarose (Apex Bioresearch Products). The kidneys were sectioned with 30μm thickness on Leica VT1000 S vibratome in PBS at room temperature. The tissue sections were stored in 1x PBS with 0.05% sodium azide. The tissue sections were rinsed with PBS. Antigen retrieval was performed by incubating tissue sections in a glass test tube containing 10mM sodium citrate with 0.05 Tween-20 solution at 80°C. After letting the sections cool at room temperature, they were rinsed thoroughly with PBS. The sections were blocked for 30mins at room temperature in blocking buffer (20mM Tris-HCl, pH 7.4, 150 mM NaCl, 200mM glycine, 0.1% v/v Tween 20, 1.0% w/v BSA, and 1.0% v/v fish gelatin). The sections were incubated at room temperature with respective primary (LC3 (1:200), #14600-1-AP from Proteintech, Rosemont, IL, USA and αSMA (1:500) # MA5-11547, from Invitrogen, Carlsbad, CA, USA) followed by secondary antibodies (Donkey anti-Rabbit cy3 #711-165-152 and Donkey anti-mouse 488 #715-545-150 from Jackson Immunoresearch, Pennsylvania, USA,) and Hoechst Live Cell Staining 33342 (#62249 from Fisher Scientific). The tissue sections were rinsed and mounted using Prolong Gold mounting medium and allowed to cure at room temperature. The immunostained section was imaged on Nikon CSU-W1 SoRa super-resolution microscope. Images were deconvolved using Nikon NIS Elements Analysis software.

**Transmission Electron Microscopy:**

Kidneys were fixed and processed as described (5) with 2.5% glutaraldehyde in 0.1 mol/L phosphate buffer (pH 7.4), followed by 1% $OsO_4$. After dehydration, thin sections were stained with uranyl acetate and lead citrate. Images were taken digitally under a JEOL JEM-1400 Transmission Electron Microscope

**Quantification of cyst:**

The kidney tissue sections (5µm) were stained with Hematoxylin and Eosin and images were taken using Nikon 80i upright microscope (Tokyo, Japan), and cyst number, % cystic index (total cyst area/ total kidney area) were quantified using ImageJ (Fiji, Madison, WI, USA) (6).

**Quantitative real-time PCR:**

RNA was isolated from mouse kidney tissue or cultured cells using trizol method (#15596018, Invitrogen, Carlsbad, CA, USA). High-capacity cDNA reverse transcription kit from Applied Biosystems (# 4368814, Foster City, CA, USA) was used to make cDNA according to the manufacturer’s protocol. Quantitative real-time PCR was done using power SYBR Green PCR master mix from Applied Biosystems (#A25742 Foster City, CA, USA) according to the manufacturer’s protocol. The mouse and human primer list are provided in the Supplementary Table 1a and 1b respectively. 18S mRNA levels were measured to normalize gene expression. The individual Ct values obtained from RTPCR reaction cycle was used to calculate 2ˆ(–delta delta Ct). The control groups (WT or RC/RC) were normalized to 1 and the fold-change was represented as a graph for individual genes.

**References:**

1. Jamadar A, Dwivedi N, Mathew S, Calvet JP, Thomas SM, and Rao R. Vasopressin Receptor Type-2 Mediated Signaling in Renal Cell Carcinoma Stimulates Stromal Fibroblast Activation. *International Journal of Molecular Sciences.* 2022;23(14):7601.

2. Norregaard R, Tao S, Nilsson L, Woodgett JR, Kakade V, Yu AS, et al. Glycogen synthase kinase 3alpha regulates urine concentrating mechanism in mice. *Am J Physiol Renal Physiol.* 2015;308(6):F650–60.
3. Sinha S, Dwivedi N, Woodgett J, Tao S, Howard C, Fields TA, et al. Glycogen synthase kinase-3beta inhibits tubular regeneration in acute kidney injury by a FoxM1-dependent mechanism. *FASEB J.* 2020;34(10):13597–608.
4. Singh SP, Tao S, Fields TA, Webb S, Harris RC, and Rao R. Glycogen synthase kinase-3 inhibition attenuates fibroblast activation and development of fibrosis following renal ischemia-reperfusion in mice. *Dis Model Mech.* 2015;8(8):931–40.
5. Abrahamson DR, and St John PL. Loss of laminin epitopes during glomerular basement membrane assembly in developing mouse kidneys. *J Histochem Cytochem.* 1992;40(12):1943–53.
6. Jamadar A, Ward CJ, Remadevi V, Varghese MM, Pabla NS, Gumz ML, et al. Circadian Clock Disruption and Growth of Kidney Cysts in Autosomal Dominant Polycystic Kidney Disease. *J Am Soc Nephrol.* 2024.

**Supplemental Table-1**

**Table 1A: Mouse primers sequence used or QRTPCR**

| No | Gene Name | Primer Sequence |
|---|---|---|
| 1 | αSMA *(Acta2)* | F – TCAGGGAGTAATGGTTGGAATG<br>R - GGTGATGATGCCGTGTTCTA |
| 2 | *Col1a1* | F - AGACATGTTCAGCTTTGTGGAC<br>R - GCAGCTGACTTCAGGGATG |
| 3 | *Col3a1* | F - TCCCCTGGAATCTGTGAATC<br>R - TGAGTCGAATTGGGGAGAAT |
| 4 | *Col5a1* | F - AGATGGCATCCGAGGTCTGAAG<br>R - GACCTTCAGGACCATCTTCTCC |
| 5 | Fibronectin (*Fn1*) | F - ATGTGGACCCCTCCTGATAGT<br>R - GCCCAGTGATTTCAGCAAAGG |
| 6 | 18S | F - GTAACCCGTTGAACCCCATT<br>R - CCATCCAATCGGTAGTAGCG |

**Table 1B: Human primers sequence used or QRTPCR**

| No | Gene Name | Primer Sequence |
|---|---|---|
| 1 | *BNIP3* | F - TCAGCATGAGGAACACGAGCGT<br>R - GAGGTTGTCAGACGCCTTCCAA |
| 2 | *BECN1* | F - CTGGACACTCAGCTCAACGTCA<br>R - CTCTAGTGCCAGCTCCTTTAGC |
| 3 | *SESN1* | F - TCACAGTGTGGATGAGATGCCG<br>R - CTCGACATTCCTGTAACTGCCTC |
| 4 | *BCL-2* | F - ATCGCCCTGTGGATGACTGAGT<br>R - GCCAGGAGAAATCAAACAGAGGC |
| 5 | *BCL-XL* | F - ATGTGGACCCCTCCTGATAGT<br>R – GCCCAGTGATTTCAGCAAAGG |
| 6 | *18S* | F - ACCGCGGTTCTATTTTGTTG<br>R - CCCTCTTAATCATGGCCTCA |
| 7 | *COL1A1* | F- AGACATGTTCAGCTTTGTGGAC<br>R- GCAGCTGACTTCAGGGATG |
| 8 | *VIM* | F- AGGCAAAGCAGGAGTCCACTGA<br>R- ATCTGGCGTTCCAGGGACTCAT |
| 9 | *αSMA (ACTA2)* | F - TCAGGGAGTAATGGTTGGAATG<br>R - GGTGATGATGCCGTGTTCTA |
| 10 | *COL3A1* | F- TGG TCT GCA AGG AAT GCC TGGA<br>R- TCT TTC CCT GGG ACA CCA TCAG |
| 11 | *COL5A1* | F- GGAGATGATGGTCCCAAAGGCA<br>R- CCATCATCTCCTTTGTCACCAGG |

| | | |
|---|---|---|
| 12 | *FN1* | F-ATGTGGACCCCTCCTGATAGT<br>R -GCCCAGTGATTTCAGCAAAGG |
| 13 | *FSP1* | F-CAGAACTAAAGGAGCTGCTGACC<br>R-CTTGGAAGTCCACCTCGTTGTC |
