## Supplementary material for "Myofibroblast- specific autophagy drives cyst growth in autosomal dominant polycystic kidney disease": Unedited western blot images

Figure 1D

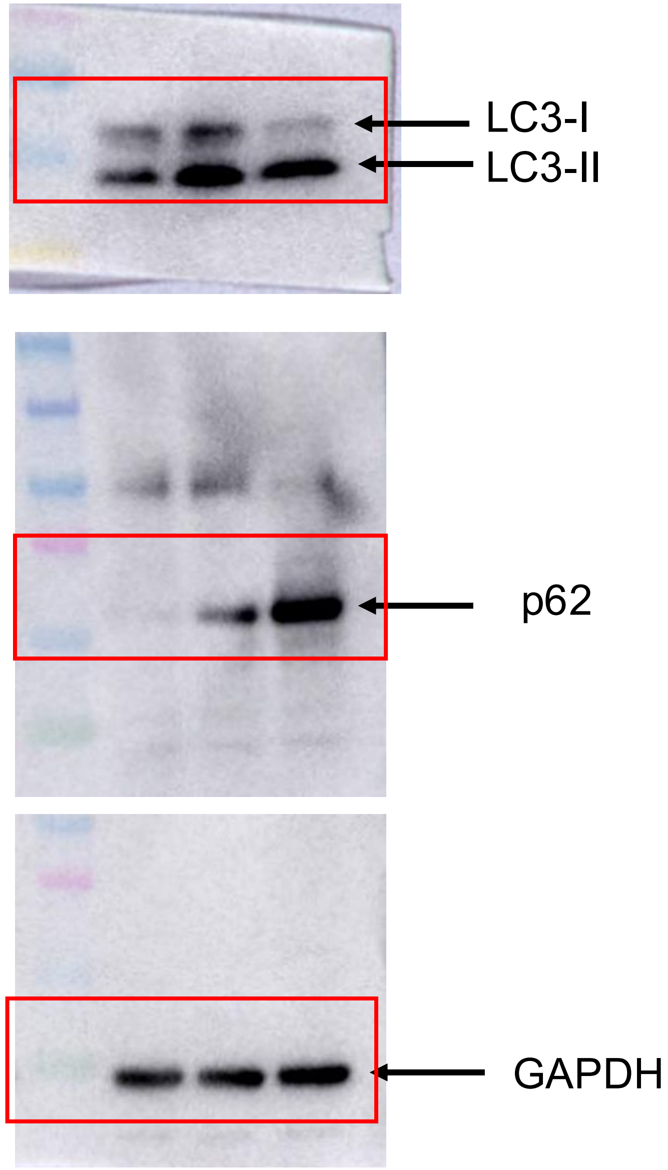

Figure 3 A

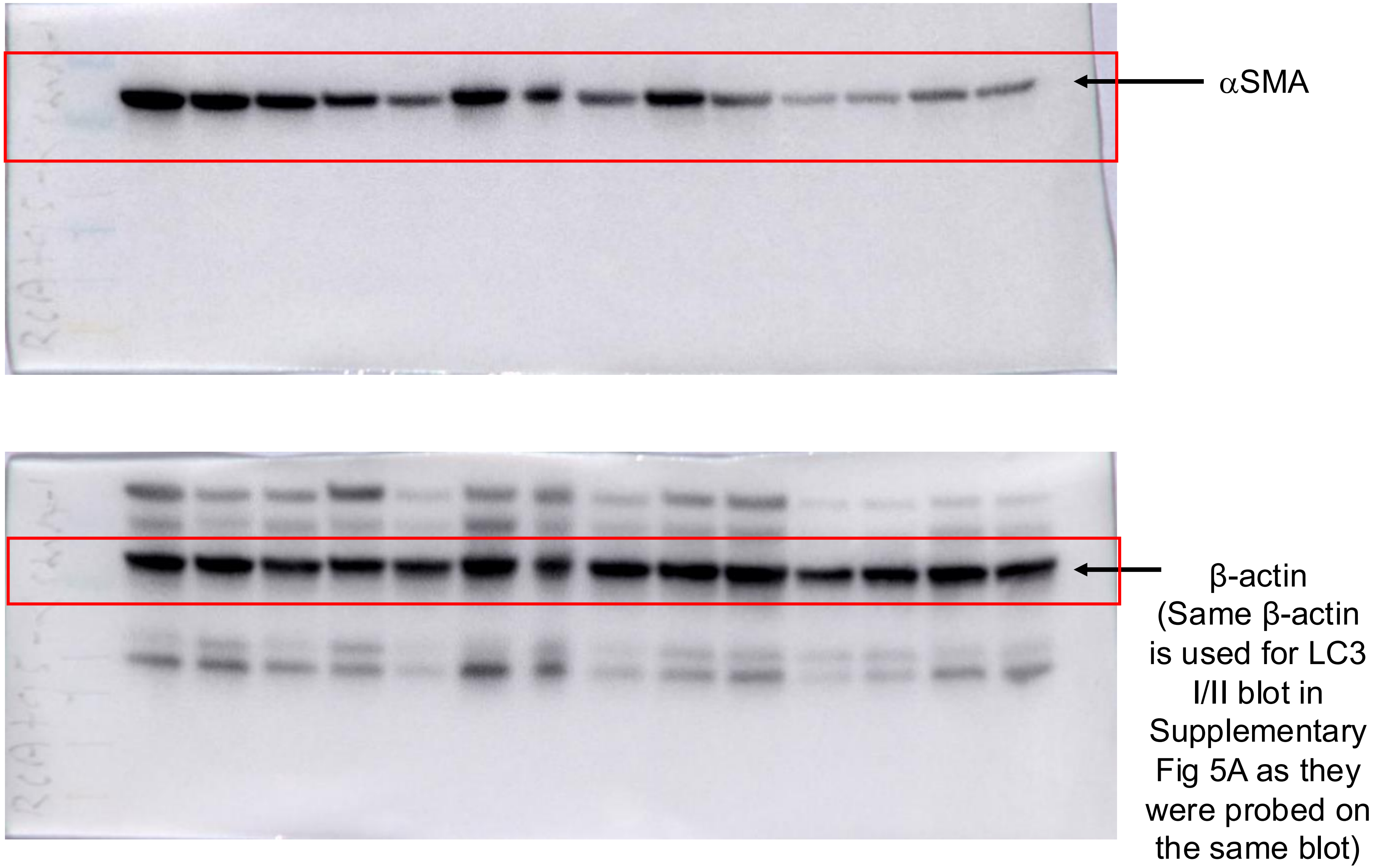
αSMA
β-actin
(Same β-actin
is used for LC3
I/II blot in
Supplementary
Fig 5A as they
were probed on
the same blot)

Figure 4 C

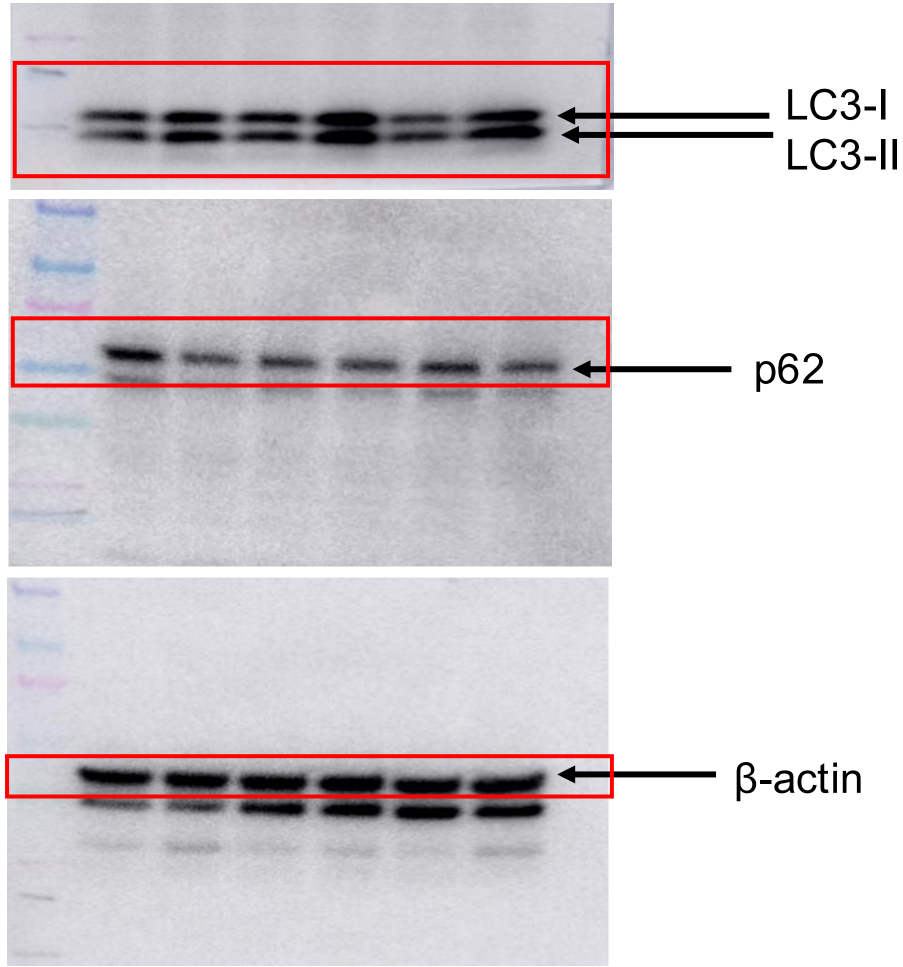


Figure 4 F

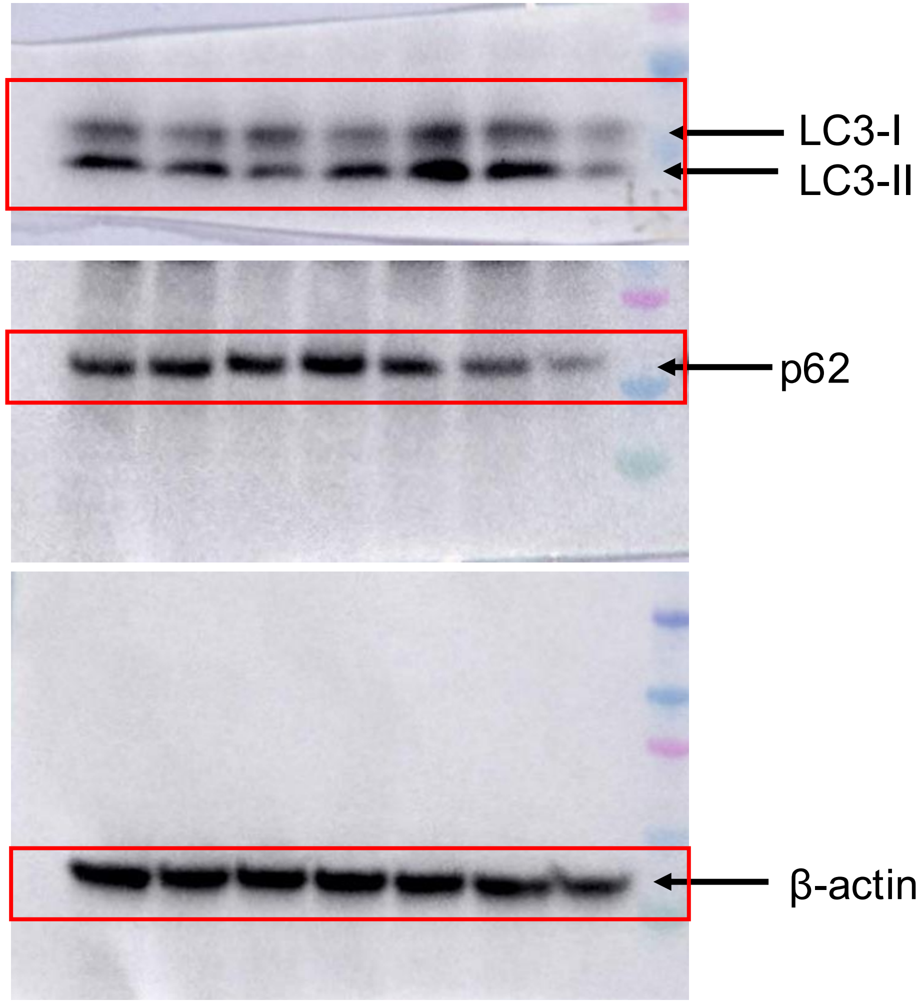

Figure 5B

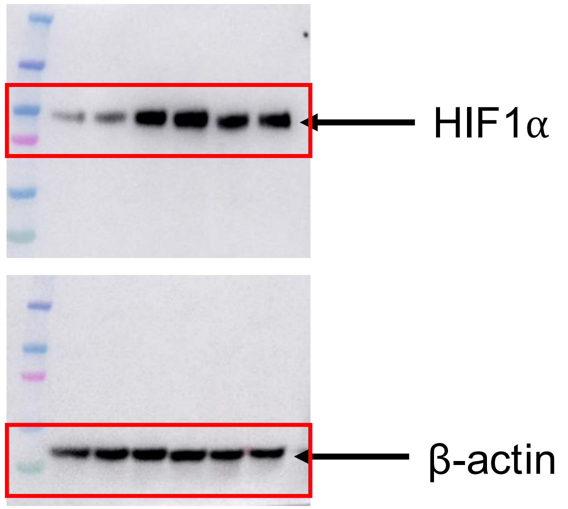


Figure 5D

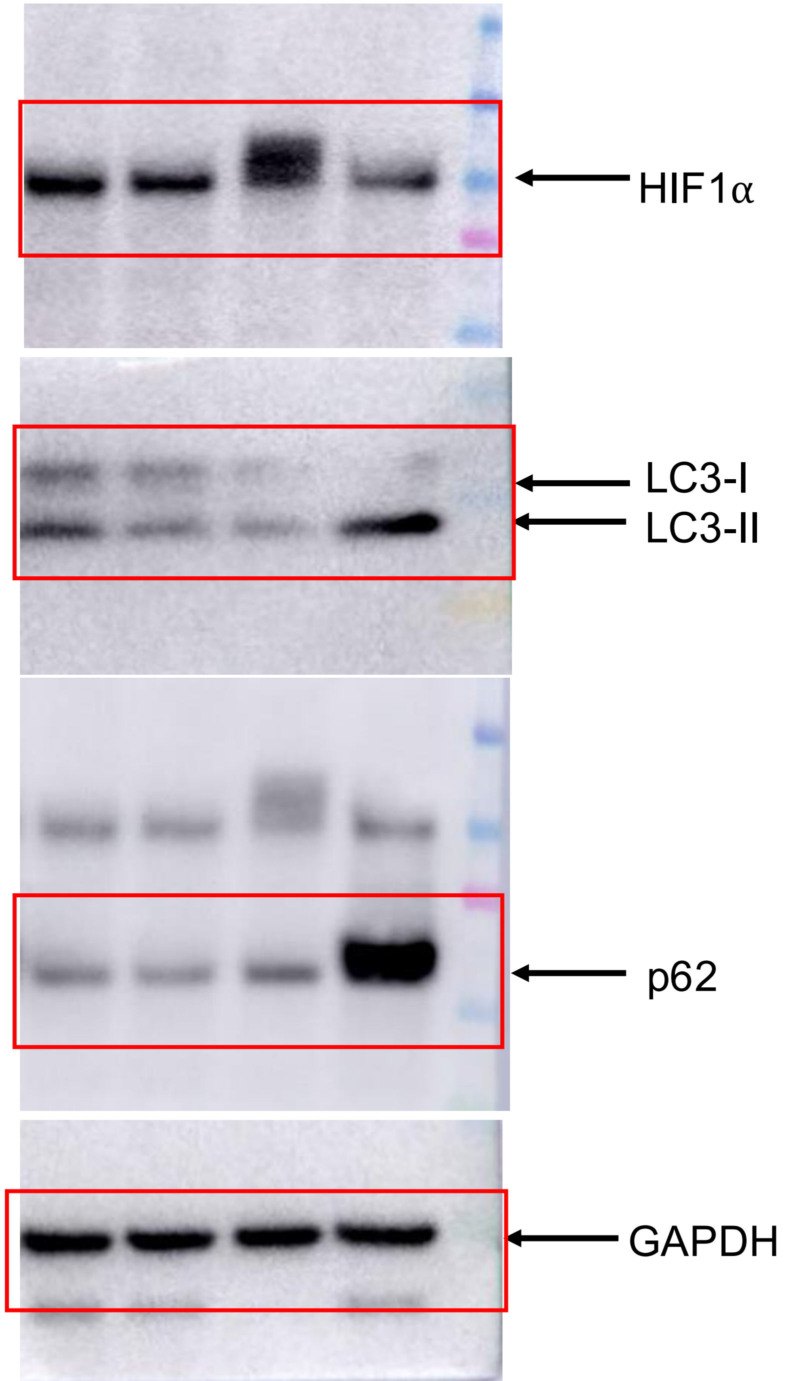

Supplementary Figure 4

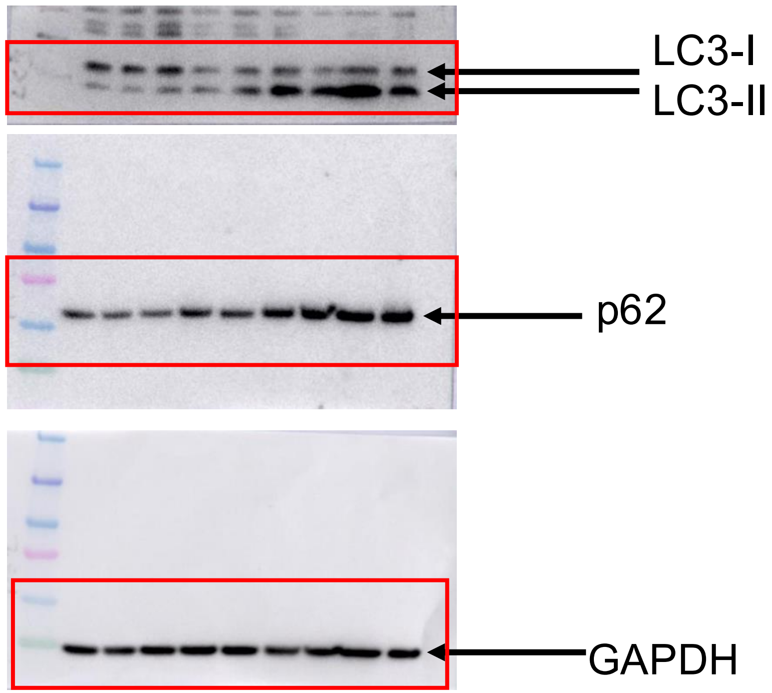

Supplementary Figure 5A

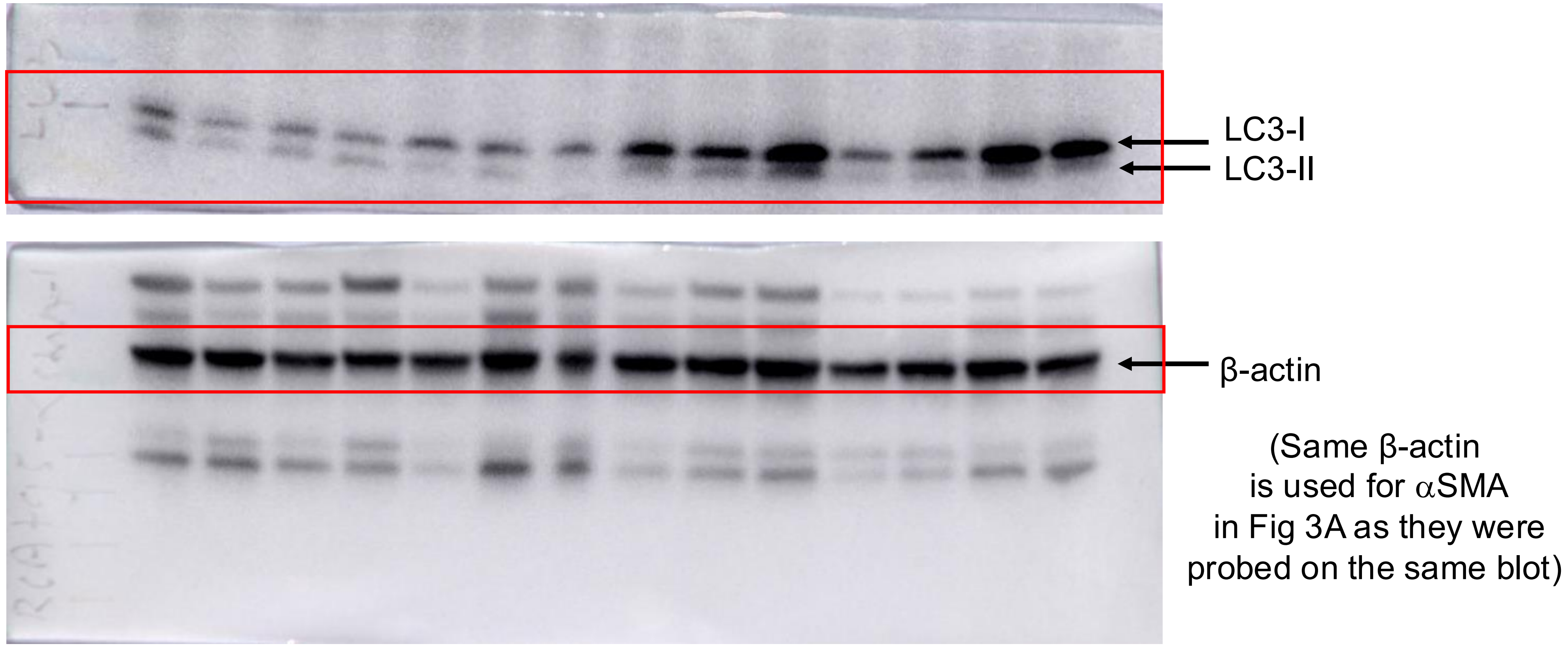

Supplementary Figure 5A

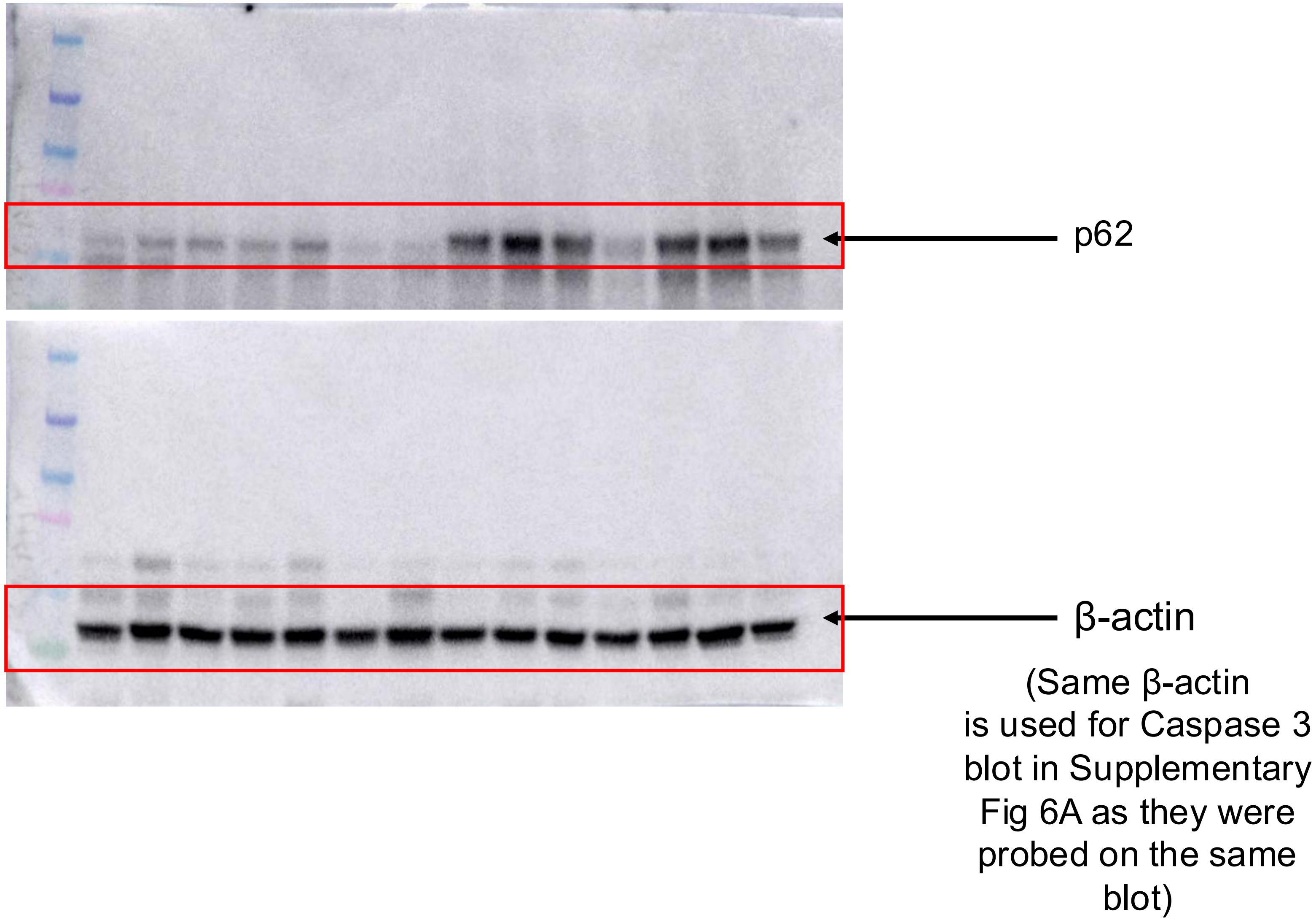

Supplementary Figure 6A

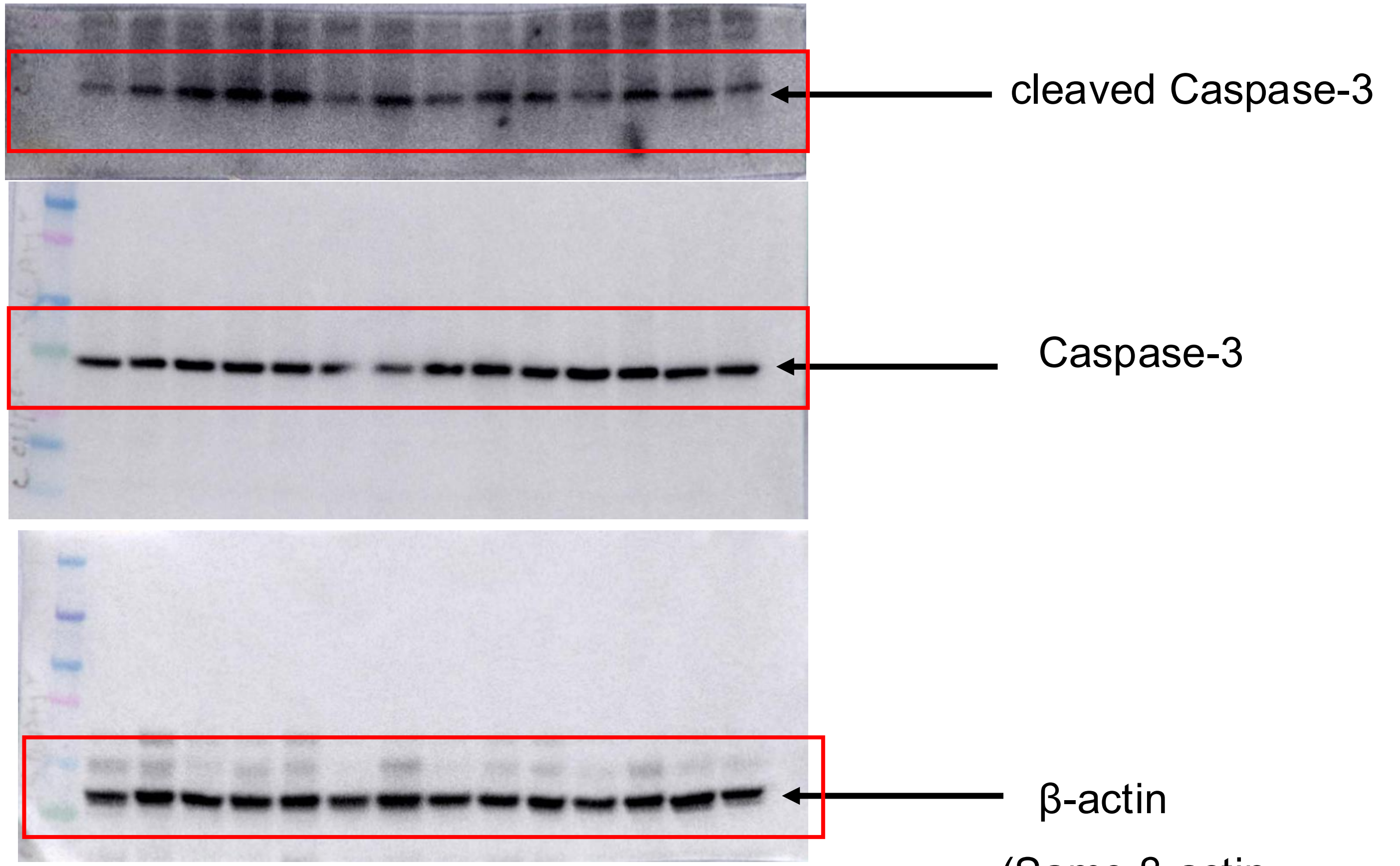


(Same β-actin is used for p62 in Supplementary Fig 5A as they were probed on the same blot)

Supplementary Figure 8A

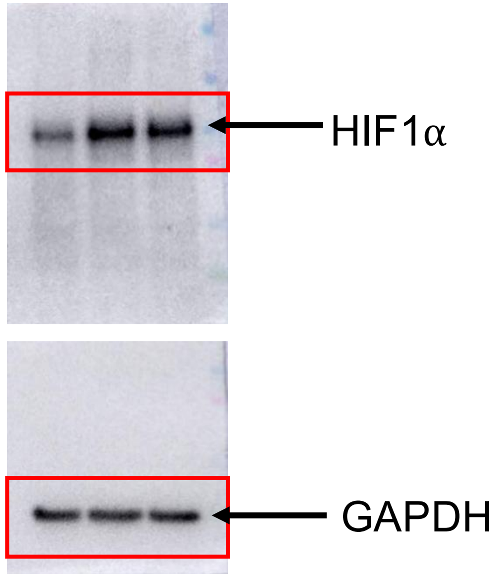


Supplementary Figure 8B

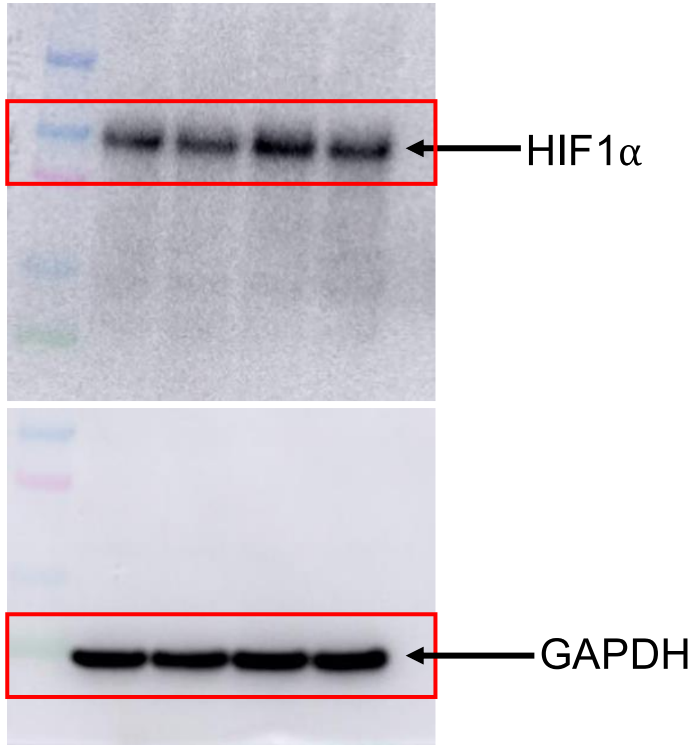

### Supplementary Figure 8C

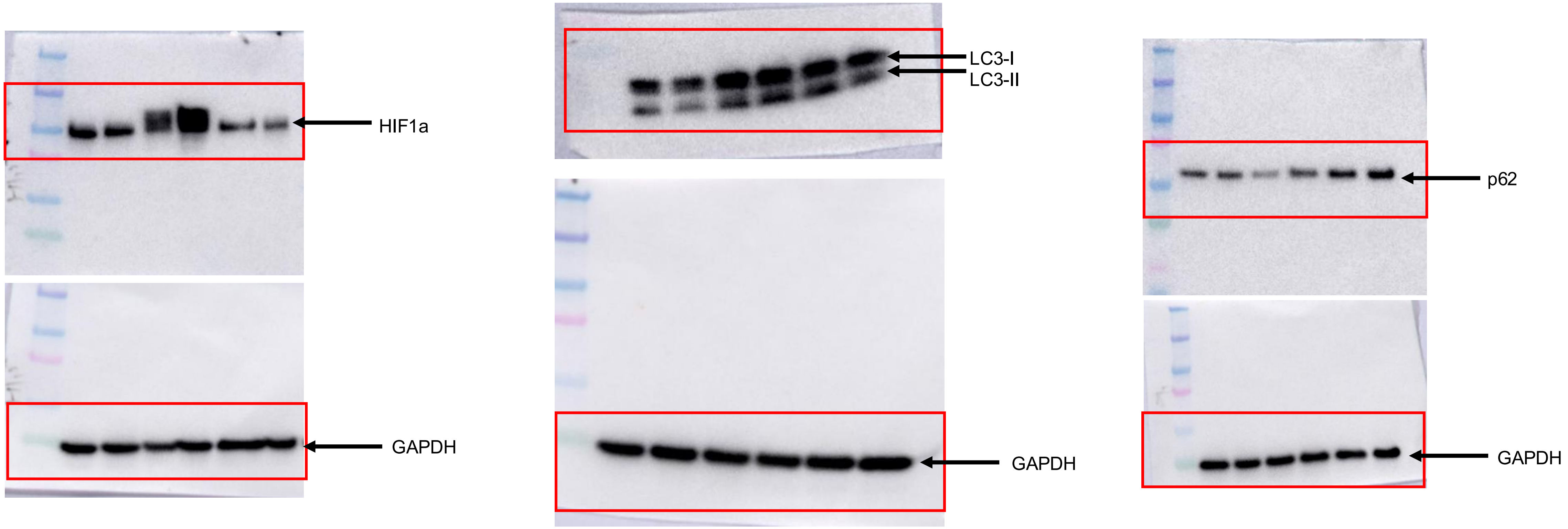
